# MYC modulates TOP2A diffusion to promote substrate detection and activity

**DOI:** 10.1101/2025.07.15.665031

**Authors:** Donald P Cameron, Kathryn Jackson, Alessia Loffreda, Carl Möller, Vladislav Kuzin, Matteo Mazzocca, Evanthia Iliopoulou, Evgeniya Pavlova, Bea Jagodic, Brian Saidel Lopez Duran, Valérie Lamour, Fredrik Westerlund, Davide Mazza, Laura Baranello

## Abstract

Topoisomerases alleviate DNA supercoiling by cleaving and resealing DNA strands. Previously, we showed that the oncoprotein MYC recruits and stimulates topoisomerases to remove DNA entanglements generated by oncogenic transcription. Understanding this mechanism may suggest methods to inhibit MYC-driven topoisomerase activation, targeting tumor-specific transcription. Here, we demonstrate that the essential topoisomerase TOP2A in human cells exists in a dynamic equilibrium between sequestration in the nucleolus, substrate searching in transcription hubs, and active engagement on chromatin. This equilibrium is highly responsive to changes in DNA topology, allowing cells to regulate TOP2A levels. Using single molecule tracking, we show that MYC accelerates TOP2A diffusion in cells. We explain this phenotype by demonstrating that MYC limits TOP2A self-interaction *in vitro*, while decreasing the size of TOP2A complexes in cells. By increasing TOP2A diffusion, MYC promotes substrate binding and increases TOP2A engagement on chromatin genome-wide, revealing the mechanism underlying MYC stimulation of TOP2A activity.

## Introduction

MYC functions as a global activator of transcription, increasing the amplitude of transcriptional bursts and boosting promoter output^1^. Transcription also generates DNA supercoiling downstream and upstream of an elongating RNA Polymerase (RNAP)^2^. As MYC increases transcription rates, it drives more torsional stress into the DNA template which, if not resolved, can slow down transcription and hinder MYC-driven overexpression^3^. Thus, MYC’s ability to increase transcriptional output depends on efficient relief of topological problems.

Topoisomerases relax DNA supercoils and decatenate intertwined DNA by cutting and resealing DNA strands. Topoisomerase 1 (TOP1) creates a transient single-strand break, allowing rotation around the unbroken strand to release DNA twist. Topoisomerase 2 (TOP2) generates a transient double-stranded break to pass one intact DNA duplex through the break, thus resolving supercoils or DNA catenanes. Both enzymes facilitate transcription by relieving topological stress. Although topoisomerases exhibit limited sequence specificity, their activity is directed by recognition of DNA topology or interaction with partner proteins^4^. Previously, we demonstrated that MYC can recruit and stimulate both TOP1 and TOP2A, localizing them to sites of high transcription, thereby promoting increased transcription rates^5^. However, the precise mechanism by which MYC mediates topoisomerase recruitment and stimulation remains unclear.

TOP2A, the isoform associated with proliferating cells, has a disordered carboxyl-terminal domain (CTD) that mediates both chromatin targeting and catalytic function^6,7^. Notably, TOP2A can form CTD-dependent phase-separated condensates *in vitro*, which partition with plasmid DNA at high concentrations^8^. These condensates reflect a transition in the activity of TOP2A from relaxation and decatenation to catenation, suggesting that TOP2A phase-separates on chromatin during mitosis to induce chromatid catenation, which is essential for chromosome condensation^9^. During interphase, transcription condensates containing MYC form at active sites, including super-enhancers, to drive transcription activation^10^. Because these condensates are often associated with elevated transcription^10,11^, they are likely to include TOP2A to relieve topological stress. Although TOP2A condensates that favor catenation may be beneficial for mitotic condensation, this action would hinder the removal of topological stress generated by transcription in interphase, if local levels of TOP2A reach concentrations associated with condensate formation. It remains unclear whether TOP2A supports transcription via association with the interphase condensates.

In this study, we demonstrate that TOP2A maintains a dynamic equilibrium between nucleolar and transcription condensates in human cells. This equilibrium is maintained by the integrity of these condensates and is highly responsive to changes in DNA supercoiling. We found that MYC accelerates TOP2A diffusion in cells. In line with biophysical principles, MYC reduces the size of the TOP2A condensates *in vitro* and decreases the size of TOP2A-containing complexes in cells, providing a rationale for the enhanced diffusion. By increasing TOP2A diffusion, MYC favors TOP2A substrate detection and promotes TOP2A catalytic engagement on the chromatin genome-wide, revealing the mechanism for MYC stimulation of TOP2A activity.

## Results

### TOP2A exists in an equilibrium between nucleolar and transcription condensates

We first examined TOP2A localization in interphase HCT116 cells, a colorectal cancer cell line previously used for studies of topoisomerases^5,12^. We applied stimulated emission depletion (STED) microscopy^13^ to achieve sub-diffractive optical resolution of TOP2A. Co-staining of nucleolar substructures—the fibrillar center (RNAPI), the dense fibrillar compartment (Fibrillarin), and the granular component (Nucleophosmin)—revealed that TOP2A was enriched across all three compartments (Fig. 1a and Extended Data Fig. 1a). Since rRNA transcription occurs specifically at the boundary of the fibrillar center^14^, this indicates that the nucleolar TOP2A enrichment is not specific to its known site of activity during ribosomal RNA (rRNA) transcription^15^. Following treatment with buffer to remove soluble proteins^16^, TOP2A enrichment in the nucleolus persisted, but was lost along with nucleolar protein fibrillarin upon further treatment with RNase A (Fig. 1b and Extended Data Fig. 1d), suggesting its association with nucleolar structures was maintained by ribosomal RNA. Outside of the nucleolus, STED imaging also revealed heterogeneity of TOP2A showing enrichment in nucleoplasmic puncta (Fig. 1a and Extended Data Fig. 1b). We predicted that these puncta correspond to transcription condensates given TOP2A’s involvement in transcriptional regulation^17^. Indeed, co-staining with MED4, a component of the Mediator complex enriched in transcription condensates^18^, showed strong colocalization with TOP2A (Fig. 1c and Extended Data Fig. 1b). This was further supported by overlapping ChIP-seq signals of TOP2A and MED26—another subunit of Mediator—from previously published datasets^5,19^ at MED26 peaks (Extended Data Fig. 1c).

**Fig. 1.**
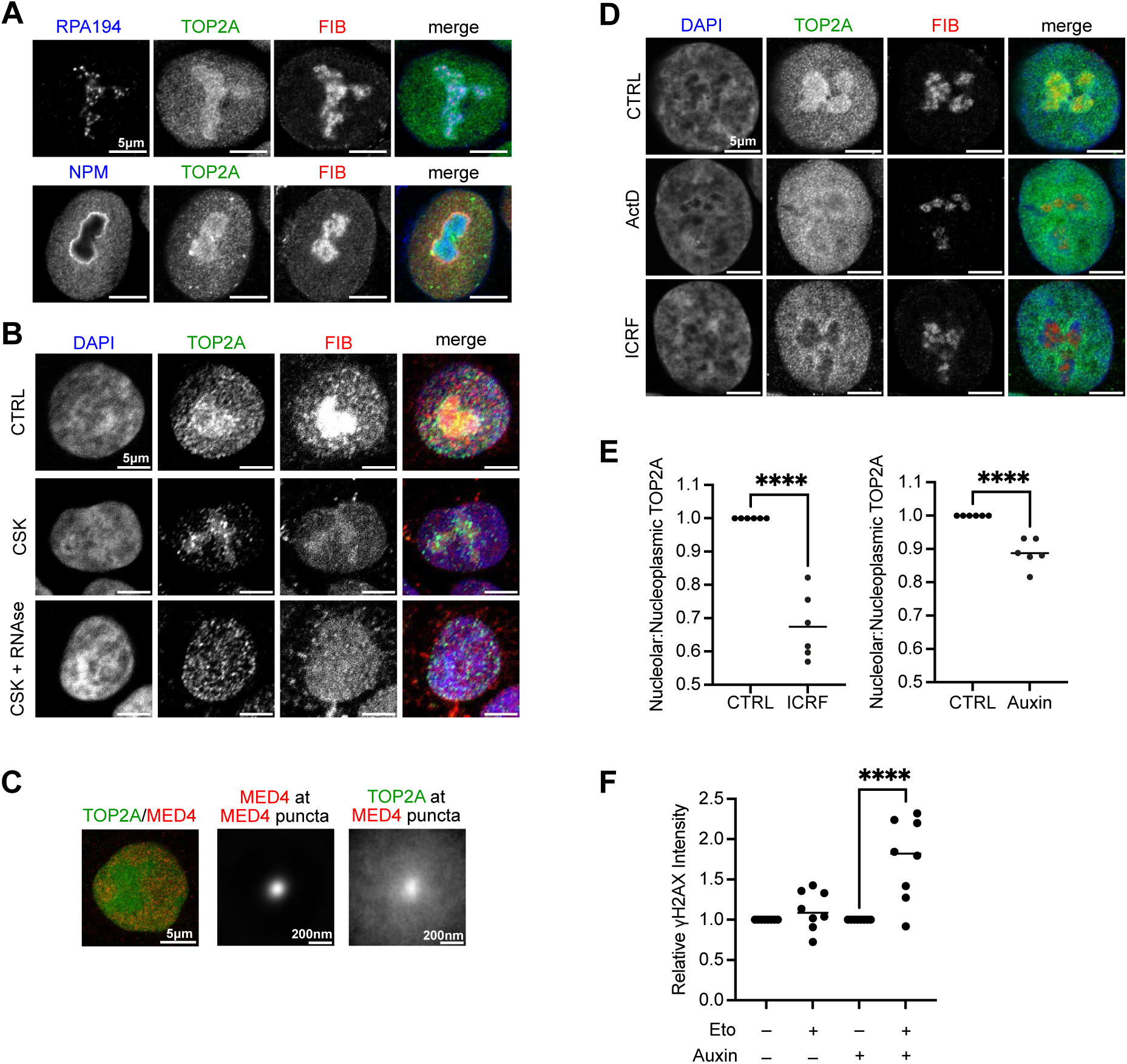
TOP2A exists in an equilibrium between nucleolar and transcription condensates. **a**, STED images of HCT116 cells immunostained for TOP2A, RNAPI subunit RPA194, nucleolar proteins Fibrillarin (FIB) and Nucleophosmin (NPM). Two independent experiments were performed. Scale bar = 5 μm. **b**, Confocal images of HCT116 cells treated with CSK buffer to remove soluble proteins +/− RNAse immunostained for TOP2A and FIB, and DNA stained with DAPI. Four independent experiments were performed. Scale bar = 5 μm. **c**, Representative STED image of HCT116 cell (left) immunostained for TOP2A and Mediator component MED4. Average MED4 (middle) and TOP2A (right) signal at 8,240 MED4 puncta from 30 cells from three independent experiments. Scale bar as indicated. **d**, STED images of HCT116 cells treated with TOP2 inhibitor 5 μM ICRF-193 for 30 minutes or 5 nM Actinomycin D (ActD) for 1 hour to specifically inhibit RNAPI, immunostained for TOP2A, and FIB, and DNA stained with DAPI. Two independent experiments were performed. Scale bar = 5 μm. **e,** Quantitation of nucleolar:nucleoplasmic ratio of TOP2A signal normalized to untreated control (CTRL) after treating HCT116 cells (left) with 5 μM ICRF-193 for 15 minutes or HCT116^TOP1-AID^ cells (right) with 500 μΜ auxin for 1 hour, to degrade TOP1. Six independent experiments were performed. 550 CTRL cells or 604 ICRF-treated cells (left) and 732 CTRL cells or 889 auxin-treated cells (right) were measured. ****p < 0.0001 (unpaired t test). **f**, Nuclear γH2A.X intensity of HCT116^TOP1-AID^ cells +/− 500 μM auxin for 1 hour +/– 1 μM etoposide (Eto) treatment for 1 hour, normalized to respective “– Eto” conditions. Eight independent experiments were performed. 1020 CTRL cells, 1085 Eto-treated cells, 1188 auxin-treated cells, or 1378 auxin and Eto-treated cells, were measured. ****p < 0.0001 (Ordinary one-way ANOVA, Šidák correction).

Previous studies have demonstrated that TOP2A exists in an equilibrium between the nucleolus and the nucleoplasm^20,21^. Given the enrichment of TOP2A in the nucleolus and nucleoplasmic transcription condensates, we propose that the integrity of these compartments supports this equilibrium. To challenge this hypothesis, we first observed how TOP2A distribution changes after these two compartments are targeted by drug treatment. Inhibition of RNAPI elongation with 5 nM actinomycin D for 1 hour reduced ribosomal RNA expression (Extended Data Fig. 1e) and induced nucleolar-to-nucleoplasmic translocation of TOP2A (Fig. 1d and Extended Data Fig. 1f), as previously reported with other RNAPI transcription inhibitors^21^. Given that this treatment did not extensively affect the nucleolar structure as observed by fibrillarin staining (Fig. 1d), it suggests that TOP2A is largely retained in the nucleolus via rRNA-mediated nucleolar interactions. Conversely, treatment with 1,6-hexanediol—shown to rapidly dissolve Mediator-containing transcriptional condensates^18^—caused TOP2A to shift from the nucleoplasm into the nucleolus within 5 minutes (Extended Data Fig. 1g). While 1,6-hexanediol can dissolve many condensates by disrupting weak hydrophobic interactions, it can favor condensation of other compartments through hydrophilic interaction^22^. Indeed, other phase separated proteins that are maintained through hydrophilic interactions are not dissolved by 1,6-hexanediol treatment^23^. This could include nucleolar proteins such as nucleophosmin, which is enriched in the nucleolus by its positively and negatively charged disordered domain^24^. This may explain why TOP2A enrichment in the nucleolus is not disrupted by 1,6-hexanediol treatment. Since the dysregulation of either the nucleolus or nucleoplasmic transcription condensates causes the rapid translocation of TOP2A from one region to the other, the data suggest the equilibrium is at least in part maintained by these two structures.

The presence of a dynamic equilibrium might allow for rapid TOP2A translocation upon changes in DNA topology. We predicted that the TOP2A equilibrium enables the cell to rapidly respond to changes in the supercoiling burden. If this is the case, increasing the level of supercoiling in the genome should provoke a redistribution of TOP2A from the nucleolus to the nucleoplasm. We tested this hypothesis in two ways: i) knowing that TOP1 and TOP2 functions in the cells are partially compensatory, we acutely degraded TOP1 by auxin, and ii) we treated cells with ICRF-193, which blocks TOP2A recycling after DNA religation^25,26^. Both treatments have been reported to increase genomic supercoiling by psoralen binding assay^27^. In both tested conditions, we observed a rapid translocation of TOP2A from the nucleolus to the nucleoplasm and nucleolar periphery (Fig. 1d,e and Extended Data Fig. 2a,b). To test whether the translocated TOP2A was actively engaged on the DNA, we treated cells with a low dose (1 μM) of etoposide for 60 minutes. Etoposide traps covalently engaged TOP2 on the DNA, causing DNA damage, which we can visualize by γ-H2AX staining. We observed an increase in DNA damage only after TOP1 degradation by auxin and etoposide treatment, suggesting that the increased nucleoplasmic fraction of TOP2A after TOP1 degradation correlates with increased TOP2 activity (Fig. 1f and Extended Data Fig. 2c). These γ-H2AX puncta were enriched within the nucleoplasm relative to the nucleoli, indicating that the increased activity is spread across the genome rather than being specific to the ribosomal DNA loci (Extended Data Fig. 2d,e). It is also possible that part of the γ-H2AX signal is associated with the pre-existing nucleoplasmic TOP2A trying to compensate for the absence of TOP1. These findings suggest that TOP2A localization is regulated by its interaction with both nucleolar and transcription condensates, existing in an equilibrium between compartments that is sensitive to both their dysregulation and DNA supercoiling.

Notably, TOP2B distribution across the nucleus was much more homogeneous as compared to TOP2A, although ICRF-193 also caused a slight re-localization nucleolus-to-nucleoplasm (Extended Data Fig. 2f,g) suggesting it may also be responsive to similar stimuli to TOP2A, although the TOP2A response was more pronounced.

### The dynamics of TOP2A in the nucleus can be described by three states

To better characterize the dynamic equilibrium of TOP2A between its associated compartments, we tagged endogenous TOP2A with a Halo-tag and used a custom Highly Inclined Laminated Optical sheet (HILO) imaging platform^28^ to perform Single Molecule Tracking (SMT) in live HCT116 cells. This approach allowed us to measure the location and motion of single fluorescently labeled TOP2A molecules over time (Fig. 2a). Single TOP2A molecules exhibited a wide range of frame-to-frame displacements, and their mobility was substantially reduced when TOP2A was trapped on DNA by ICRF-193 treatment^29^ (Fig. 2b).

**Fig. 2.**
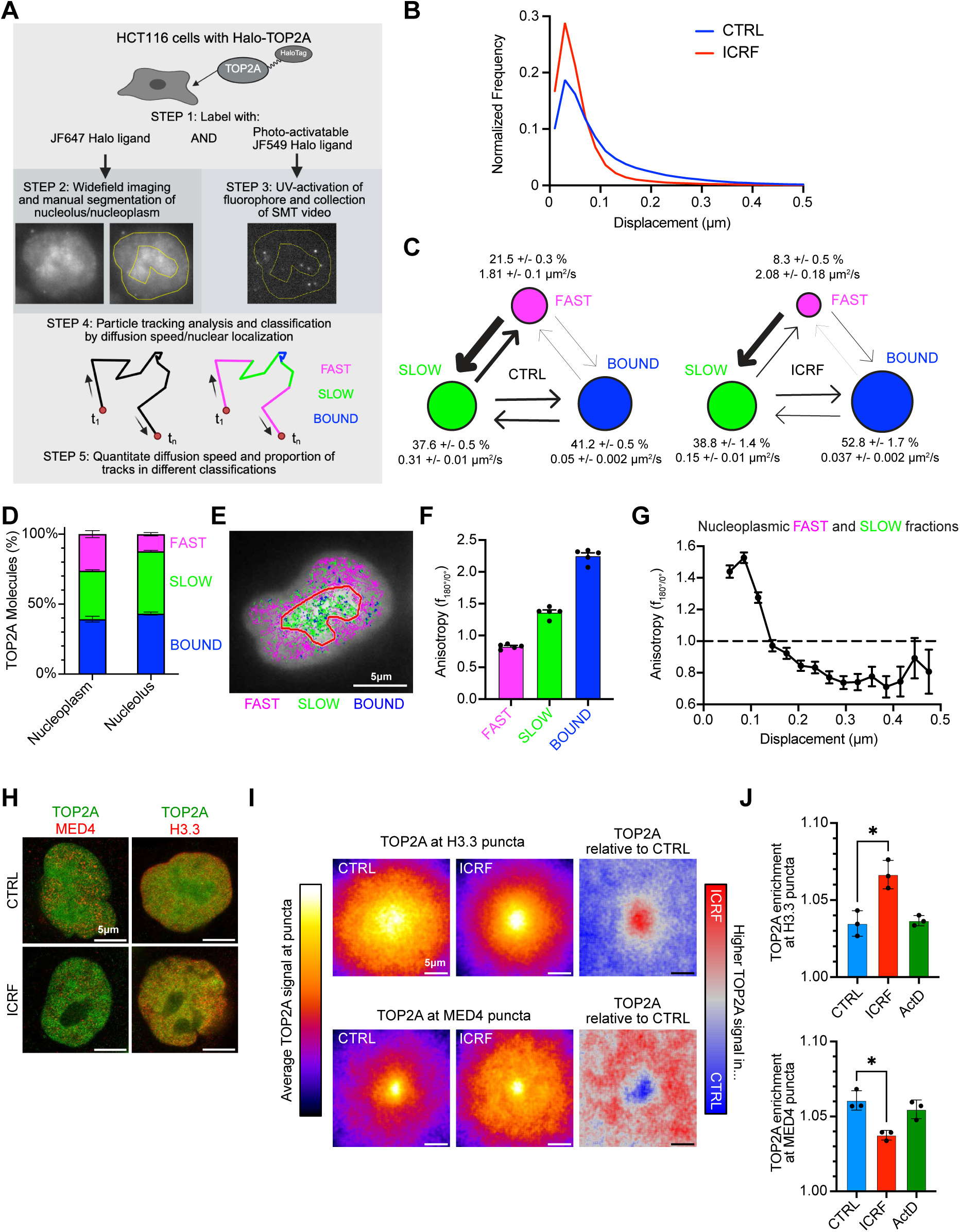
TOP2A exists in various diffusive states within different compartments of the nucleus. **a**, Scheme of SMT detection and analysis. **b**, Representative histogram of linear distance traveled by TOP2A molecules between 10ms frames from SMT in Halo-TOP2A HCT116 cells +/− 5 μΜ ICRF-193 for 15 minutes. Histogram normalized to maximum bin value of each experiment. **c**, Percentage of TOP2A in Fast, Slow and Bound fractions and their relative diffusion coefficient in control conditions and upon 5 μΜ ICRF-193 for 15 minutes. Width of arrows is proportional to frequency of detected transitions between states. Average of 4 independent experiments, 12 image fields per condition per experiment. **d**, Percentage of Fast, Slow and Bound fractions within nucleoplasm and nucleolus. **e**, Representative image of Halo-TOP2A HCT116 cell stained for TOP2A (white) with nucleolus circled in red, and Fast, Slow and Bound TOP2A tracks highlighted in pink, green and blue, respectively. Nucleolus is identified by GFP-Nucleolin transfection and detected through Structured Illumination light sheet microscopy (SIM). **f**, Average anisotropy values of Fast, Slow and Bound tracks from Halo-TOP2A HCT116 cells. 5 independent experiments, 115 cells total. **g**, Anisotropy values (f180/0) vs mean displacement (μm) of combined Fast and Slow tracks from Halo-TOP2A HCT116 cells. **h**, Representative STED images of HCT116 cells labelled with TOP2A and either MED4 or H3.3 +/− 5 μM ICRF-193 treatment for 30 minutes. **i**, Average TOP2A signal at H3.3 (top) and MED4 (bottom) puncta from STED images after control (left) or 5 μM ICRF-193 treatment for 30 minutes (center), and difference between both conditions (right). Signal averaged from three independent experiments. 7177-14486 puncta per condition. **j**, Average TOP2A puncta enrichment across independent experiments from data in Figures 2I and S3A. *p < 0.05 (ANOVA, Dunnett’s multiple comparison test).

Using a Hidden Markov Model approach (variational Bayes single particle tracking, vbSPT)^30^, we found that nuclear TOP2A diffusion could be categorized into three distinct populations. The slowest population had diffusion coefficients below 0.1 µm^2^/s (Fig. 2c), comparable to the mobility of histones and chromatin bound proteins^31^. We refer to this group as the bound fraction. The other two populations, which represent TOP2A molecules diffusing in the nucleus, are referred to as slow and fast fractions based on their diffusion rates (Fig. 2c). Treatment with ICRF-193, which prevents TOP2A disassociation from DNA after one cycle of catalytic activity^25,26^, caused a marked increase in the bound population. As it takes over an order of magnitude longer for strand passage by TOP2A to occur than the framerate of our imaging (300 ms for strand passage^32,33^ compared with 10 ms between frames), and since DNA-bound TOP2A exhibits diffusion rates far lower than TOP2A in the fast and slow fraction, we conclude that this fraction includes TOP2A covalently engaged on the DNA. However, this may also include TOP2A that is interacting with chromatin without catalytic engagement. TOP2A in the fast and slow states is not precluded from becoming covalently engaged on the DNA, given the observed transitions between the fast and slow fractions as well as between the slow and bound fractions (see arrows in Fig. 2c). It is notable that fast-to-slow and slow-to-bound transitions are substantially more frequent than fast-bound transitions, suggesting the TOP2A molecules must pass through the slow state before becoming chromatin-bound.

Using a GFP-Nucleolin marker to separately analyze TOP2A diffusion in the nucleolus and the nucleoplasm, we observed that diffusion was reduced by more than 50% in the nucleolus relative to the nucleoplasm with a rate of 0.61 μm^2^/s in the nucleoplasm as compared to 0.28 μm^2^/s in the nucleolus. In addition, the nucleolus predominantly contained slow and bound TOP2A, while the nucleoplasm contained more equal amounts of all three populations (Fig. 2d,e). This indicates that TOP2A diffusion is more constrained within the nucleolus. The fast and slow fractions are further distinguishable by their diffusional anisotropy, with the nucleoplasmic slow fraction exhibiting anisotropy values greater than 1 (Fig. 2f), indicative of protein retention within discrete compartments. Nucleoplasmic tracks with average anisotropy greater than 1 were limited to regions of about 150 nm in size (Fig. 2g), corresponding to both the approximate size of the MED4 puncta (Fig. 1c) and the reported size of a transcriptional condensate^34^. These features suggest that the slow TOP2A fraction in the nucleoplasm likely contains molecules performing compact target searching in transcription condensates^35^.

If the slow TOP2A fraction reflects TOP2A localized in transcription condensates and the bound fraction corresponds to TOP2A interacting with chromatin, then we should be able to visualize the TOP2A equilibrium shift upon ICRF-193 treatment given that transcription condensates transiently associate with gene loci^11^. To test this, we used STED to observe TOP2A association with transcription condensates labelled with MED4 and TOP2A association with actively transcribed chromatin labelled with H3.3, which we predict to contain TOP2A molecules belonging to the slow and bound fractions, respectively^36^ (Fig. 2h). TOP2A was enriched in both compartments, indicating overlap with transcriptional condensates and transcribed chromatin (Fig. 2i). Notably, ICRF-193 treatment shifted TOP2A from the MED4 compartment to the H3.3 compartment (Fig. 2i,j), consistent with its movement from the slow fraction to the bound fraction (Fig 2c). This shift was not observed with actinomycin D treatment, ruling out the possibility that it was driven solely by increased nucleoplasmic translocation (Fig. 2j and Extended Data Fig. 3a).

Overall, these results suggest that nucleoplasmic TOP2A can be grouped into three states based on diffusion rates: a bound fraction interacting or catalytically engaged with chromatin, a slow fraction enriched in transcription condensates, and a fast fraction diffusing through the nucleoplasm. The equilibrium maintained between these states and the nucleolus enables TOP2A to rapidly shift to regions where topological regulation is needed.

### MYC modulates TOP2A diffusion in cell

We have previously shown that the oncogenic transcription factor MYC is able to join TOP1 and TOP2A to form the TOPOisome. Within this complex, MYC stimulates the catalytic activity of both topoisomerases, thereby coupling oncogenic transcription rates to relief from DNA entanglements^5^. If the formation of a TOPOisome prevents the binding of other proteins thereby limiting the effective size of the complex—as shown in the MYC-driven TOPOisome study—then we would expect to observe differences in TOP2A diffusion with or without MYC since protein diffusion is inversely correlated with the complex radius^37^.

We generated HCT116 cells expressing MYC tagged with an auxin-inducible degron (AID)^38^, enabling 95% knockdown of MYC protein within 90 minutes of auxin treatment (Fig. 3a). We then tagged endogenous TOP2A *via* Halo-tag and performed SMT to track TOP2A diffusion upon auxin-dependent MYC degradation. Strikingly, we found that although the proportion of TOP2A found in the fast, slow, and bound fractions were largely unchanged (Fig. 3b), TOP2A diffusion speed was reduced after MYC degradation in both the fast and slow fractions of the nucleoplasm (Fig. 3c,d). However, we found no difference in the nucleolus (Fig. 3e) where MYC is largely excluded (Extended Data Fig. 4a). This reduction in TOP2A diffusion was not observed upon global transcription inhibition by triptolide treatment^39^, indicating that the diffusion change observed upon MYC depletion was not linked to changes in transcription (Extended Data Fig. 4b-d). Having confirmed that TOP2A diffuses faster in the presence of MYC, in line with our prediction that MYC limits TOP2A complex size, we wanted to test if this phenotype could be observed both *in vitro* and in cells.

**Fig. 3.**
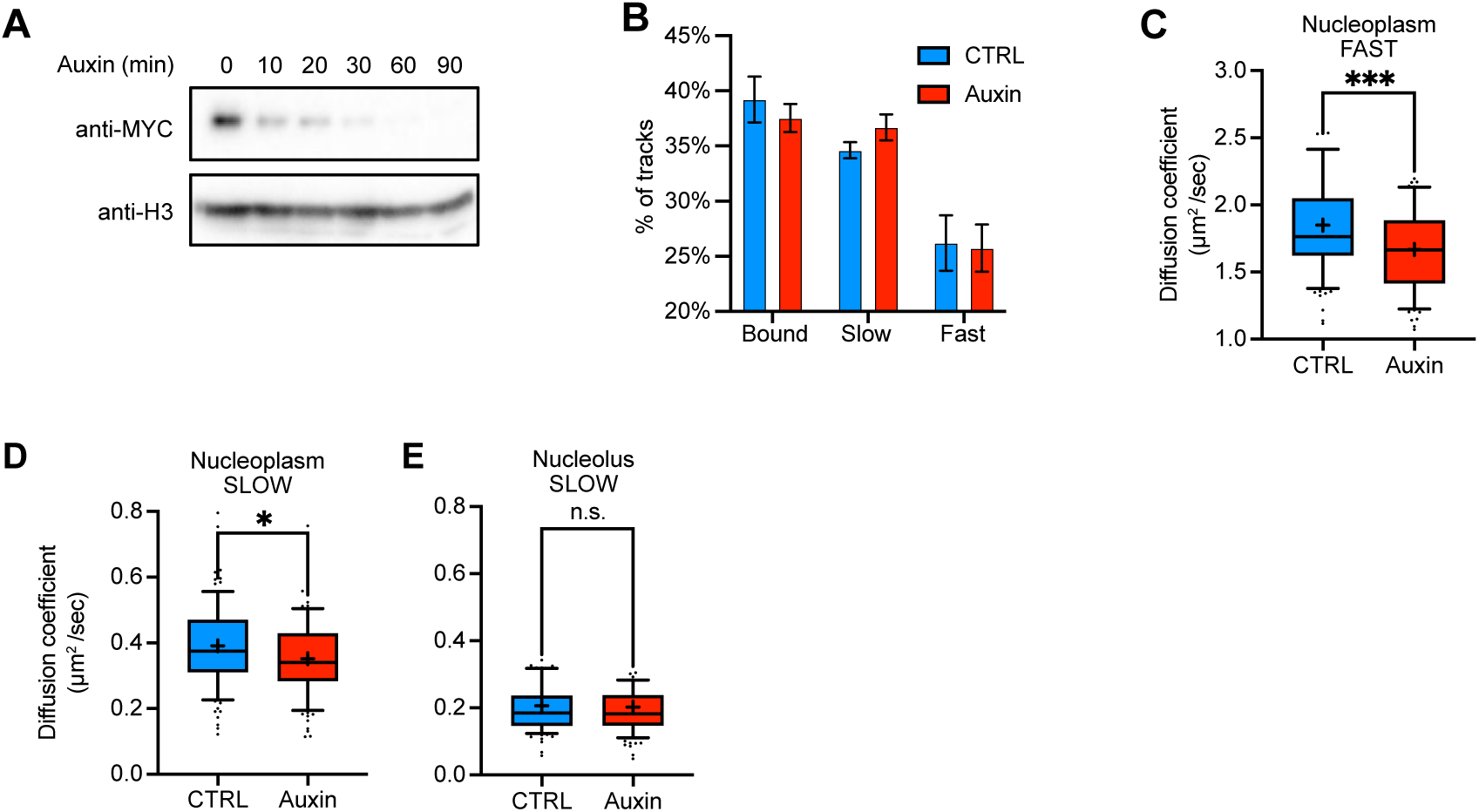
Loss of MYC reduces TOP2A diffusion in cells. **a**, Representative western blotting demonstrating near complete degradation of MYC in HCT116^MYC-AID^ cells within 90 minutes treatment with 500 μM auxin. Two independent experiments. **b**, Proportion of TOP2A in Fast, Slow and Bound fractions from HCT116^MYC-AID^ cells +/− auxin treatment. Average of five independent experiments. **c**-**e**, Diffusion coefficients of Fast nucleoplasmic (**c**), Slow nucleoplasmic (**d**), and Slow nucleolar (**e**) fractions from individual Halo-TOP2A HCT116^MYC-AID^ cells. Average of four independent experiments, 100 cells for each condition. ***p < 0.001; *p < 0.05 (Unpaired t-test).

### MYC limits TOP2A droplet size

*In vitro,* we took advantage of recent findings that TOP2A forms phase-separated droplets^8^. The droplets are largely dependent on the presence of the TOP2A-CTD, which is intrinsically disordered. We were able to observe droplet formation using 2 μM recombinant human TOP2A in an isotonic salt solution that was enhanced both by co-incubation with DNA and reduction in salt concentration (Fig. 4a-b and Extended Data Fig. 5a,b), in line with previous reports^8^. These droplets were also able to fuse (Extended Data Fig. 5c). Furthermore, we observed that while supercoiled plasmid DNA and catenated kDNA—both known substrates of TOP2A—as well as RNA could increase droplet formation, short 20 bps oligonucleotides did not (Fig. 4a-b), confirming previous data that DNA was required to be at least 50-100 bps to favor droplets formation^8^.

**Fig. 4.**
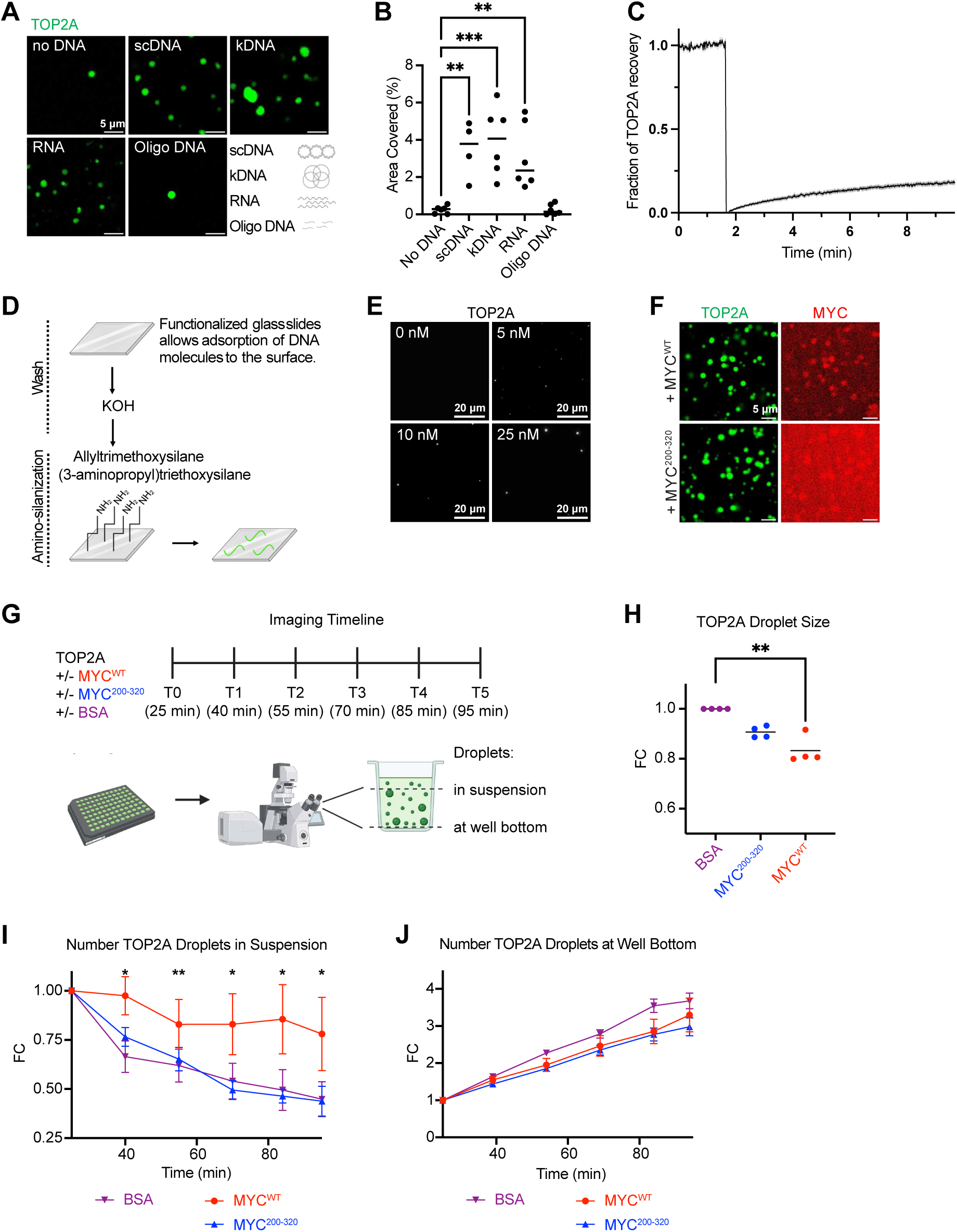
MYC affects the biophysical properties of TOP2A condensates *in vitro*. **a**, Representative images of 2 μM AF488-labelled SNAP-TOP2A condensates in the absence or presence of 10 ng/μl negatively supercoiled plasmid DNA (scDNA), catenated kinetoplast DNA (kDNA), RNA, or 20 bp oligonucleotides (oligo DNA). **b**, Quantitation of percentage of field area covered by conditions in Fig. 4a. Average of 4-6 independent experiments. ***p < 0.001; **p < 0.01 (ordinary one-way ANOVA). **c**, FRAP experiment of SNAP-TOP2A condensates demonstrating limited fluorescence recovery after droplet bleaching. Average of five independent experiments. **d**, Scheme of droplet deposition onto silanized slides used in Fig. 4e and Extended Data Fig. 5j. **e**, Visualization of TOP2A condensates after addition of scDNA deposited onto functionalized glass coverslips. Contrast differs for each image to distinguish signal from noise and demonstrate condensate formation at low concentrations. **f**, Representative images of co-incubation of AF488-labelled SNAP-TOP2A with AF647-labelled SNAP-MYC^WT^ or SNAP-MYC^200–320^. **g**, Schematic of imaging droplets over time. **h**, Normalized quantitation of 0.5 μM SNAP-TOP2A droplet size +/− 3.5 μM BSA, MYC^WT^, or MYC^200–320^ in the presence of 10ng/μM scDNA. Average of six independent experiments. **p < 0.01; *p < 0.05 (Kruskal-Wallis test Dunn’s correction). **i**,**j**, Quantitation of TOP2A droplets from 2 μM SNAP-TOP2A +/− 3.5 μM BSA, MYC^WT^ or MYC^200–320^ in suspension (**i**) or at well bottom (**j**) over the course of 95 minutes normalized to the 25 minutes time point. **p < 0.01; *p < 0.05 (multiple paired t tests).

The dependence on DNA length to stimulate droplet formation implies the existence of a percolated network structure within the droplets where physical crosslinks form between molecules such as through ionic interactions^40^. This contrasts with classical liquid-liquid phase separation (LLPS) where a density transition occurs at the saturation concentration^41^. If these droplets form through LLPS, we would expect to observe free diffusion between the dilute and droplet phases and a strict threshold for determining the concentration where phase separation occurs. However, we observed limited diffusion between phases as assessed by fluorescence recovery after photobleaching (FRAP) (Fig. 4c), in contrast to near complete fluorescence recovery as seen with liquid-liquid phase separated-droplets formed by the protein FUS^42^. To image TOP2A condensates at protein concentrations below previously reported saturation concentrations, we adapted a protocol where glass cover slips are functionalized to generate a surface with net positive charge that attracts the negatively charged DNA, thus keeping the condensates stationary^43^ (Fig. 4d). Using this technique, we could observe droplet formation in the presence of plasmid DNA at protein concentrations as low as 5 nM (Fig. 4e). The variance in puncta size indicated that these droplets are made up of multiple TOP2A molecules, excluding the possibility that we observed individual TOP2A molecules. These data suggest that TOP2A droplets form by percolation and that TOP2A condensates can still form at sub-saturating concentrations of proteins.

Since MYC can also bind TOP2A independently of TOP1^5^ (Extended Data Fig. 5d,e), we tested whether MYC can affect TOP2A condensate formation *in vitro*. Incubation of fluorescently labeled recombinant TOP2A and MYC^WT^ with plasmid DNA showed that MYC co-localizes with TOP2A droplets (Fig. 4f) albeit in a non-stoichiometric manner (Extended Data Fig. 5f). Interestingly, while the MYC mutant containing only residues 200-320 (MYC^200–320^) does not interact with TOP2A in immunoprecipitation experiments^5^, it did show weak co-localization with TOP2A droplets (Fig. 4f and Extended Data Fig. 5g). While others have reported that MYC can phase-separate in the presence of crowding agents such as PEG^10^, we did not observe MYC-only droplets under the conditions used here (Extended Data Fig. 5h).

To understand how MYC affects the properties of TOP2A droplets, we devised an imaging strategy that allowed us to observe droplets both in solution and at the well bottom (Fig. 4g). This allowed the visualization of droplets independently of interaction with the plastic surface. We combined 500 nM of fluorescently labeled TOP2A and supercoiled plasmid DNA with 3.5 μM MYC^WT^, MYC^200–320^, or BSA in an isotonic buffer and measured the resulting TOP2A droplet properties. MYC^WT^ induced a decrease in TOP2A droplet size (Fig. 4h). Similar results were seen at 2 μM (Extended Data Fig. 5i) and 25 nM (Extended Data Fig. 5j) of TOP2A. In line with the above observation, the TOP2A droplets formed in the presence of MYC^WT^ exhibited a markedly slower rate of sedimentation over a time course of 95 minutes (Fig. 4i,j) as demonstrated by comparing the relative number of droplets that remain in suspension *vs* those at the well bottom. This is in contrast to co-incubation of TOP2A with MYC^200–320^ where sedimentation rates were consistent with the BSA control, suggesting that co-localization of MYC^WT^ within TOP2A condensates also reduces their droplet density.

We noted that MYC^200–320^ elicited a limited decrease in TOP2A droplet size (Fig. 4h and Extended Data Fig. 5i), indicating that it was able to modulate TOP2A droplet formation to some degree while lacking the region required for direct interaction with TOP2A. However, these experiments were done with much higher concentration of MYC as compared to pull down conditions (100 nM) where we demonstrated that MYC^200–320^ did not bind TOP2A during immunoprecipitation. We propose this could be due to the presence of MYC^200–320^, albeit lower than MYC^WT^, in the TOP2A droplets (Fig. 4f). Thus, reducing protein concentration should abrogate this effect. Using the glass coverslip system described previously (Fig. 4d), we demonstrated that at lower protein concentration (25 nM TOP2A and 100 nM MYC), MYC^WT^ reduced the size of TOP2A droplets whereas this effect was negligible with MYC^200–320^. This suggests that the effect of the mutant required much higher concentration than MYC^WT^ (Extended Data Fig. 5j).

Note that, despite MYC’s ability to interfere with TOP2A droplet formation, MYC-TOP2A interaction does not directly depend on TOP2A’s CTD as shown by the immunoprecipitation of MYC with the TOP2A^ΔCTD^ mutant that lacks this domain (Extended Data Fig. 5k,l). Together, these data suggest that MYC reduces the size of TOP2A droplets, potentially through weakening the interactions of TOP2A with other TOP2A molecules.

### MYC limits TOP2A interactions in cells

To test whether MYC regulates the size of TOP2A-containing complexes in cells, we used K562^MYC-AID^ cells^38^, previously used to study topoisomerases upon MYC acute depletion^5^, to rapidly degrade MYC and test whether it affects the ability of TOP2A to establish interaction with other molecules. MYC was almost completely degraded upon 90-minute treatment with auxin, while TOP2A protein levels were unaffected (Extended Data Fig. 6a). Glycerol gradient centrifugations of benzonase-digested protein lysates revealed that acute MYC-depletion caused a shift of TOP2A towards higher molecular-weight complexes (Fig. 5a,b and Extended Data Fig. 6b-d). Altogether these findings suggest a novel mechanism of MYC target regulation, where the presence of MYC is required to accelerate diffusion of TOP2A by limiting the effective size of TOP2A-forming complexes, both outside (fast fraction) and within (slow fraction) transcription condensates.

**Fig. 5.**
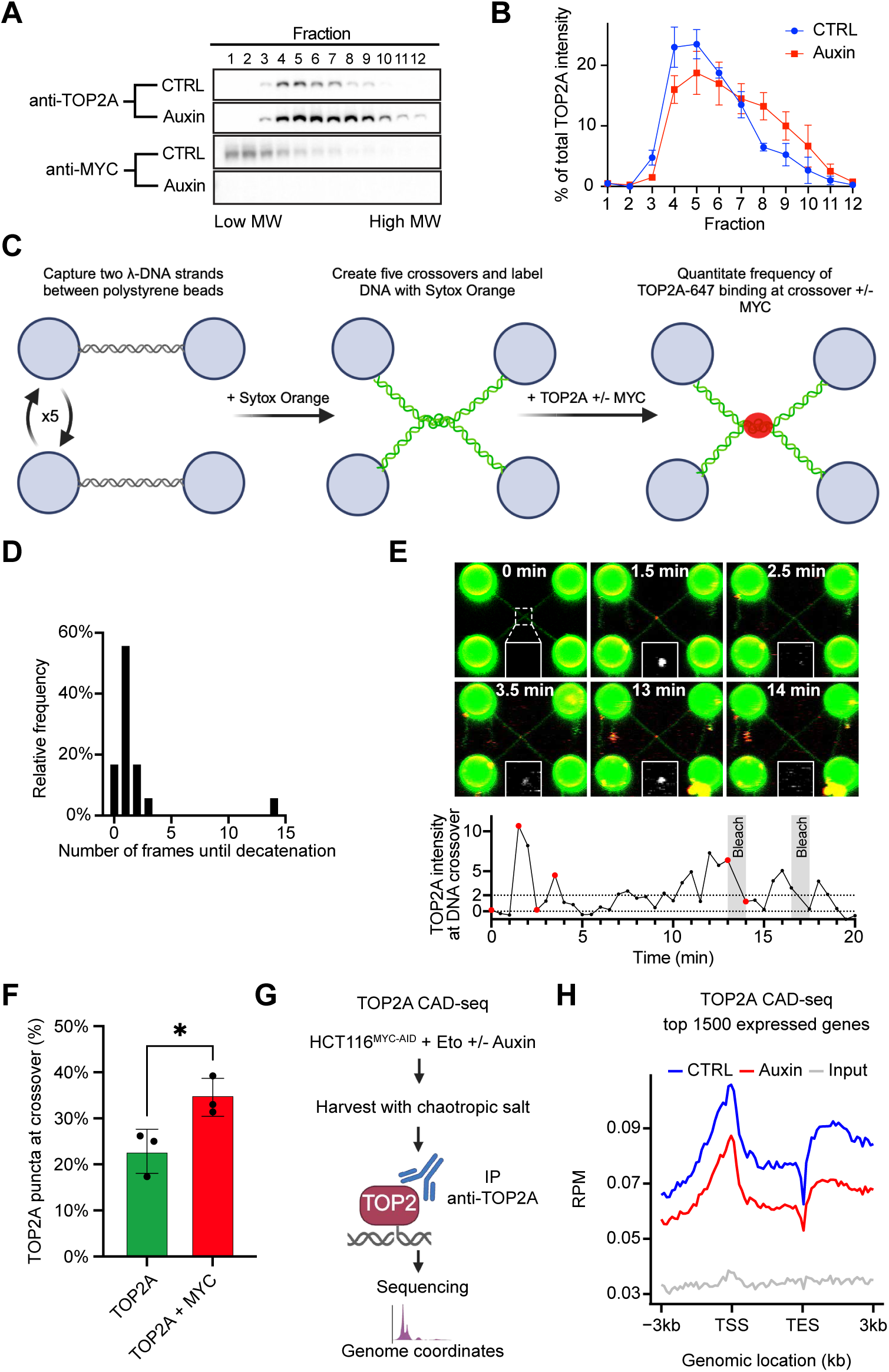
MYC limits TOP2A complex size favoring substrate detection and TOP2A activity. **a**, Representative glycerol gradient from K562^MYC-AID^ cells +/− 500 μΜ auxin treatment for 90 minutes. **b**, Quantitation of four independent glycerol gradient experiments from (A). Band intensity is normalized as a proportion of the fraction with maximum intensity to control for variation of total protein amounts. **c**, Schematic of Q-TRAP experiment. **d**, Number of frames (imaged every 4 seconds) that the DNA crossover persists after detection of TOP2A at crossover from Q-TRAP experiment in the presence of ATP (value = 0 when no TOP2A could be visualized before crossover resolution). Two independent experiments, 8-10 crossover measured per experiment. **e**, Example of TOP2A intensity measurement for DNA crossover after incubation in the absence of ATP. TOP2A intensity at crossover is normalized to background signal and quantitated every 30 seconds. TOP2A that persisted at crossover for more than 90 seconds was bleached to enable visualization of new TOP2A molecules binding to crossover. Snapshots (top) are from time points highlighted in red on graph (bottom). BSA is included in all conditions to ensure protein concentration remains consistent. **f**, Proportion of images where TOP2A is bound at the DNA crossover in the absence of ATP +/− MYC. Three independent experiments, 3-6 crossover measured for each condition per experiment. *p < 0.05 (Paired t-test). **g**, Scheme of TOP2A CAD-seq. **h**, Metagene plot of TOP2Acc enrichment across the top 1500 (∼10%) expressed genes based on published RNA-seq^12^ in HCT116^MYC-AID^ cells +/− auxin. Input DNA used as negative control. Average of three independent replicas. RPM refers to reads per million. TSS refers to the transcription start site and TES refers to the transcription end site. See individual heatmaps in Extended Data Fig. 6c.

### MYC promotes substrate detection for TOP2A

Intuitively, increasing enzyme diffusion rates should enhance substrate detection and therefore enzyme activity. We directly visualized this using quadruple trap optical tweezers (Q-TRAP) coupled with fluorescent imaging to monitor TOP2A binding at a DNA crossover formed by intertwining two linearized lambda DNA molecules (Fig. 5c). Using this system, we are also able to visualize the product of TOP2A activity by observing the decatenation of the DNA strands. Under physiological conditions including Mg^2+^ and ATP, TOP2A-mediated decatenation occurs extremely quickly (within the ∼4 seconds it takes to image another frame) such that we observe decatenation in more than 72% of instances immediately upon TOP2A binding or without even detecting the bound intermediate (Fig. 5d). The rapid nature of the decatenation reaction, combined with signal-to-noise constraints that arise when further lowering TOP2A concentration, makes it challenging to directly compare decatenation rates with or without MYC in real time using this system.

To circumvent this issue, we removed ATP so that decatenation could not occur, thereby prolonging the observation window. We then collected images at 30-second intervals and measured TOP2A localization at the crossover by applying an intensity threshold (Fig. 5e). Under these conditions, we detected more frequent TOP2A binding events when MYC was present (Fig. 5f). Given that increased substrate binding would logically translate into higher catalytic turnover when ATP is available, and, considering our prior demonstration that MYC stimulates TOP2A activity in standard assays^5^, these data strongly suggest that the MYC-enhanced TOP2A diffusion promotes greater substrate engagement, thereby increasing enzymatic activity.

### MYC increases TOP2A activity in cells

If MYC promotes TOP2A enzymatic activity by favoring substrate detection, decreasing MYC levels should decrease TOP2A-DNA cleavage complexes (TOP2Accs) in cells. To detect TOP2Accs genome-wide, we optimized our previously developed CAD-seq^44^ for TOP2A to immunoprecipitate only catalytically engaged TOP2A. HCT116^MYC-AID^ cells were treated sequentially with auxin, proteasome inhibitor MG132, and—in the last 6 minutes—etoposide to trap and stabilize TOP2ccs^45,46^ (Fig. 5g). MG132 was included to prevent TOP2cc degradation^46^. In control cells, TOP2Acc accumulated at transcription start sites (TSS) and downstream of transcription end sites (TES) of the top 1500 (∼10%) expressed genes (Fig. 5g-h and Extended Data Fig. 6e) as previously observed^47^. With MYC degradation, TOP2Accs were markedly reduced across all gene locations, particularly towards the TESs, indicating that TOP2A catalytic engagement on the DNA is MYC-dependent. This was not due to a general downregulation of transcription as 90 minutes of auxin treatment caused essentially no change in transcripts production as assessed by EU incorporation (Extended Data Fig. 6f-h). Altogether, these findings demonstrate that MYC limits the size of TOP2A complexes, increasing diffusion and substrate detection, which affects TOP2A catalytic engagement on the DNA (Fig. 6). Without ruling out the possibility of additional mechanisms, we suggest that these results reveal the primary mechanism by which MYC stimulates TOP2A.

**Fig. 6.**
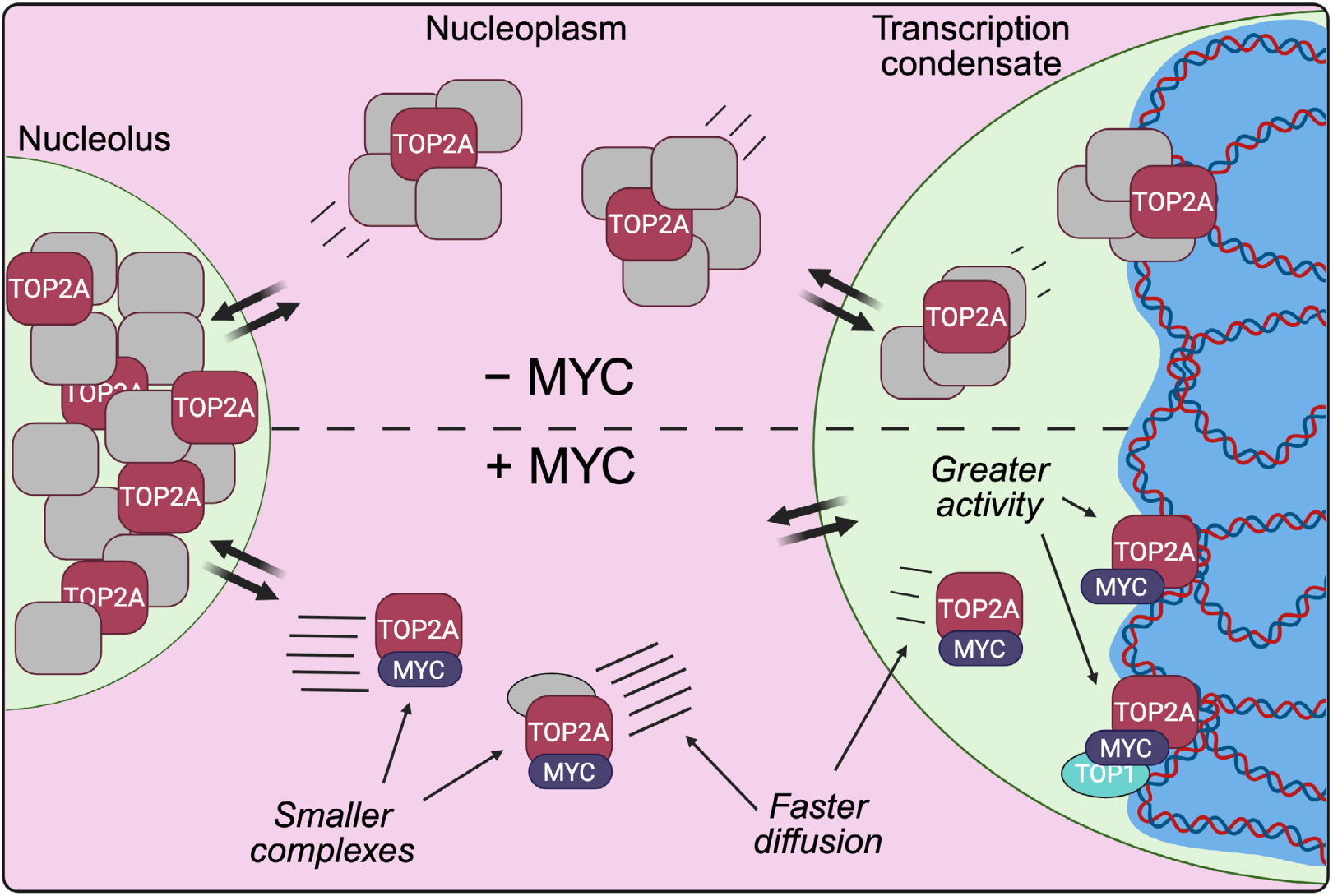
Model: MYC acts as an accelerant of TOP2A in the nucleoplasm. TOP2A exists in an equilibrium between the nucleolus, nucleoplasm, and transcription condensates, which is highly responsive to changes in supercoiling. The diffusion of TOP2A can be categorised into fast (pink), slow (green) or bound (blue), and is dependent on the cellular compartment. In the presence of MYC, TOP2A forms protein complexes, including the TOPOisome with TOP1, that are, on average, smaller than in the absence of MYC by limiting interaction with other proteins (gray boxes). This causes TOP2A to diffuse faster and increases its chromatin binding, thus stimulating its activity.

## Discussion

### The advantages of equilibrium biology

We demonstrate that TOP2A exists in a multipartite equilibrium maintained by distinct nuclear condensates and is acutely sensitive to dysregulation of topoisomerase activity (Figs. 1 and 2). There are many potential advantages to this equilibrium. For instance, TOP2A levels must be strictly regulated throughout the cell cycle to prevent aberrant activity. Its expression increases during S-phase, peaks in mitosis, and is subsequently degraded^48,49^. This ensures sufficient TOP2A is available during mitosis where it is required to condense and disentangle chromatin^50–52^, while rapid degradation prevents toxicity from excess cellular TOP2A during interphase^53^. Thus, the cell must require a mechanism to sequester excess TOP2A when expression is upregulated during G1/S-phase while also facilitating rapid degradation upon mitotic exit.

We find that TOP2A is sequestered away from nucleoplasmic DNA into the nucleolus through an RNA-dependent mechanism (Fig. 1). Previous work showed that residues 1192-1289 in the TOP2A CTD are required for RNA binding and nucleolar enrichment^54^. This region also confers sensitivity to RNA-mediated activity inhibition^54^, suggesting that RNA-rich compartments, such as the nucleolus, may repress TOP2A activity. Additionally, the nucleolus is a hub for degradation of misfolded and excess proteins^55,56^, implying it could buffer excess TOP2A by sequestration and facilitate protein degradation after mitosis. Further experiments combining quantitative imaging with acute control of TOP2A expression would be valuable to test this hypothesis.

### Sub-diffusive cellular mobility and its relevance for the TOPOisome complex

Diffusion in the cell is far slower than would be expected by macromolecules of similar sizes in a dilute solution. Molecular dynamics simulations of crowded protein environments analogous to the nucleus indicate that reduced diffusion arises from proteins forming transient clusters or complexes with other proteins^57^. These simulations demonstrate that diffusion of an individual macromolecule decreases proportionally with the number of contacts it makes with other molecules in its environment, causing a broad range of diffusion values for the same protein^58^.

We similarly detect a wide distribution of TOP2A diffusion rates (Fig. 2b). Although we group these rates into three classes (Fig. 2c) for comparison, each class likely contains multiple TOP2A sub-states. In the bound fraction, for instance, we cannot distinguish between catalytically engaged TOP2A *vs.* TOP2A merely interacting with chromatin. Since the presence of MYC increases catalytic engagement of TOP2A on genomic DNA (Fig. 5h), one might expect an increase in TOP2A bound fraction. However, since MYC limits non-specific TOP2A self-interaction *in vitro* (Fig. 4), it may also reduce TOP2A’s non-specific chromatin interactions. These counteracting effects on TOP2A-DNA interactions may explain why we could not observe a difference in the proportion of bound TOP2A +/− MYC as measured by SMT (Fig. 3b).

Moreover, while we link the slow fraction of TOP2A to transcription condensates based on anisotropy (Fig. 2f,g) and co-localization with Mediator (Fig. 2h-j), TOP2A diffusion likely depends on numerous specific and non-specific binding partners in its microenvironment. Indeed, TOP2A interacts with hundreds of proteins^59^, as well as DNA. Advances that improve both the throughput and resolution of SMT^60^ could enable further distinction of these sub-states and help define the mechanism of TOP2A diffusion in greater detail.

We also show that TOP2A diffuses faster in the nucleoplasm in the presence of MYC (Fig. 3). This finding suggests that the average TOP2A complex size is smaller when interacting with MYC. Upon downregulation of MYC by auxin treatment, the diffusion coefficients of fast and slow TOP2A molecules decrease by approximately 10%-15%. According to the Stokes-Einstein relationship, in first approximation, diffusion coefficients would scale as the cubic root of the diffusing complex, thus corresponding to roughly 30% to 50% larger complexes upon MYC degradation. We confirm this shift by glycerol gradient fractionation (Fig. 5a-b) and by measuring TOP2A droplet size as a proxy for TOP2A self-interaction (Fig. 4d-j). These results align with our prior characterization of a size-limited TOPOisome comprising TOP2A, TOP1, and MYC which stimulates topoisomerase activity *in vitro* and in cells^5^. Recently, an alternative TOPOisome where topoisomerases are nucleated and stimulated by p53 has also been identified^61^. Given that TOP2A interacts with many proteins, there may be multiple “flavors” of TOPOisome, each conferring distinct impacts on topoisomerase activity. We hypothesize that changes in TOP2A diffusion, as shown here with MYC, partially explain these distinct activity profiles. Future studies should integrate enzymatic assays with measurements of complex size and diffusion to test the generality of this mechanism.

### MYC as an accelerant

Our data indicate that MYC functions as an “accelerant”, favoring TOP2A diffusion and thereby increasing substrate detection. For a distributive enzyme such as TOP2A, increasing the likelihood to find its preferential substrate will result in increased activity, which we detect at sites of high transcription output (Fig. 5h). Acceleration of transcription factor exchange is also associated with increased transcription output highlighting the benefit of increased diffusion rates^62^. Notably, cells with greater metastatic potential have lower cytoplasmic viscosity and higher diffusivity than those with lesser metastatic potential, implying that increased diffusion may be linked to malignancy^63,64^. MYC’s established role as a transcriptional amplifier is grounded in its ability to recruit transcription factors to their targets, but how exactly it achieves this remains unclear. Indeed, MYC has many interacting partners^65^, including other enzymes involved in RNA processing, DNA replication, and chromatin modification—all processes understood to be stimulated by MYC independently of its DNA binding functions^65–67^. Interestingly, DNA-binding-deficient MYC induces proliferation in some systems^66^, but not others^68^, raising the question of whether its transcriptional function can be separated from its accelerant role. Future studies are needed to determine whether other proteins also enhance diffusion in a manner similar to MYC and to explore how cells exploit diffusion control to regulate enzymatic function.

## Methods

### Cell lines

The human colon carcinoma cell line HCT116 (ATCC, CCL-247) and variants thereof are used throughout this manuscript. Halo-TOP2A-HCT116^MYC-AID^ cells were generated by knocking in an N-terminal Halo-tag into both TOP2A alleles of HCT116 cells expressing MYC tagged with an Auxin-Inducible Degron (AID) system^38^ (gift of Dr. Zuber, (IMP, Austria)). These cells were subsequently transduced with a BFP-labeled OsTir1 delivery vector^38^ to enable MYC degradation. All HCT116 cells were cultured in high glucose (4.5 g/L) DMEM (Thermo Fisher, 61965059) with 10% Fetal Bovine Serum (Thermo Fisher, 10270106) in a 37 °C incubator with 5% CO_2_. HCT116^TOP^^1-AID^ cells^69^ were additionally cultured with 1 μg/ml puromycin (Thermo Fisher, A11138-03) and 125 μg/ml hygromycin B (Thermo Fisher, 10687010). K562^MYC-AID^ cells (gift from Dr. J. Zuber (IMP, Austria))^38^ were cultured in RPMI 1640 medium (Thermo Fisher, 21875034) supplemented with 10% Fetal Bovine Serum (Thermo Fisher, 10270106) and 2 mM L-Glutamine (Thermo Fisher, 25030081), 1 mM sodium pyruvate (Thermo Fisher, P5770598) in a 37 °C incubator with 5% CO_2_. AID-mediated degradation was induced by treatment with 500 μΜ auxin (3-indoleacetic acid, Sigma, I3750).

### Recombinant protein expression

Recombinant human TOP2A labeled at the N-terminal with the SNAP-tag sequence^70^ was expressed from the 12-URA-B plasmid in yeast cells and purified as previously described^71^. Briefly, the plasmid was transformed into URA-deficient yeast and grown in uracil-deficient media to select for transformed cells. Yeast cultures were grown in YPLG media (1% yeast extract, 2% peptone, 2% sodium DL-lactate, 1.5% glycerol) overnight and TOP2A expression was induced by addition of 2% galactose for 6 hours. The protein lysate was extracted by cryo-milling, filtered, and TOP2A was enriched using sequential HisTrap nickel and HiTrap CP cation exchange columns before cleavage with His-tagged TEV protease overnight. The TEV and cleaved Histag were removed by repassage through a HisTrap nickel column and the SNAP-TOP2A was purified using a Superdex 200 16/60 column, concentrated by filter centrifugation, and stored at −80 °C.

For expression of TOP2A^ΔCTD^, the previously published^72^ expression vector coding for the wild-type human TOP2A was mutated by site-directed mutagenesis to introduce a STOP codon at residue 1217 using the QuikChange XL Site-Directed Mutagenesis kit (Agilent), generating the hTopo IIA Δ1217plasmid. TOP2A^ΔCTD^ was overexpressed in Baby Hamster Kidney cells (BHK21). TOP2A^ΔCTD^ was first purified on a StrepTrap HP column (Cytiva, 23151816). The protein was washed with 25 mM Hepes, 100 mM NaCl, 100 mM KCl, 1 mM MgCl_2_, 10% v/v glycerol, 2 mM DTT, pH 8 and eluted with the same buffer supplemented with 3 mM Desthiobiotin (Sigma-Aldrich, 533-48-2). The Twin-strep tag was removed by the addition of P3C (PreScission protease, Sigma-Aldrich, 27-0843-01) at 1:50 ratio (w/w) and incubated overnight at 4 °C. The cleaved protein was then loaded on a HiTrap Heparin HP column (Cytiva, 17-0407-01). Elution was performed by a single step using 25 mM Hepes, 370 mM NaCl, 370 mM KCl, 1 mM MgCl_2_, 10% v/v glycerol, 2 mM DTT, pH 8.

Recombinant human MYC^WT^ and MYC^200–320^ with N-terminal SNAP-tag were expressed in Rosetta-gami competent cells in LB media. Expression of MYC was induced with 1 mM IPTG for 4 hours, and cells were lysed by probe sonication. Protein was resuspended in buffer (500 mM NaCl, 20 mM Tris pH 8, 35 mM imidazole, 1% Triton X-100) with increasing urea (0 M then 1.6 M) before being solubilized in buffer (NaCl, 20 mM Tris pH 8, 35 mM imidazole, 1% Triton X-100,7 M urea). Lysate was centrifuged, filtered before passing through a HisTrap column (Sigma, GE17-5319-01), concentrated and dialyzed to remove urea before being stored at −80 °C.

### Droplet assays

N-terminal SNAP-tagged TOP2A was labeled by incubating with 5 μM SNAP-surface Alexa Fluor 488 (NEB, S9129S) at 37 °C for 30 minutes. Excess fluorophore was removed through dialysis for 24 hours into 500 mM KCl, 20 mM Tris pH 8, 10% glycerol, 0.5 mM TCEP buffer for TOP2A. N-terminal SNAP-tagged MYC^WT^ and MYC^200–320^ were labeled in 5 μM SNAP-surface Alexa Fluor 647 (NEB, S9136S) for 37 °C for 30 minutes before being dialyzed into 20 mM Tris pH 8, 100 mM KCl, 1 mM DTT, 1 mM EDTA, 20% glycerol. BSA samples were treated in the same manner as the MYC samples.

Droplet assays done with 500 nM or 2 μΜ TOP2A and 3.5 μΜ MYC^WT^, MYC^200–320^, or BSA were mixed with 10 ng/ul pUC19 plasmid DNA or water and allowed to settle for 25 minutes before imaging over 95 minutes. Assays done using 500 nM TOP2A were imaged in six locations per well once every 25 minutes, and images were taken at the well bottom and 6 additional z-stacks with increments of 10 μm. Assays done using 2 μM were imaged in one location per well every 5 minutes once at the bottom of the well and again focusing 25 μm higher within the well. Imaging was done using a Nikon CrEST X-Light V3 spinning disk microscope. Analysis was done using CellProfiler. Droplet assays were performed in 384 well black non-binding μCLEAR microplates (Greiner, 781906).

### Confocal imaging and immunofluorescence

HCT116 cells were plated on 18 mm coverslips (Marienfeld, 630-2200) in 12-well plates to reach 70% confluency. ICRF-193 (Enzo Life Sciences, BML-GR332) treatments were done using 5 μM for 15 minutes. Etoposide (Merck, E1383) treatments were done using 1 μM for 1 hour. Actinomycin D (Merck, A1410) treatments were done using 5 nM for 1 hour. Auxin induced degradation in HCT116^TOP1-AID^ cells was done using 500 uM auxin for 3 hours or blocked using 0.1 mM auxinole overnight. After treatments, cells were washed once with PBS before being fixed with 3% paraformaldehyde (Thermo Fisher, 28908) for 5 minutes at room temperature and permeabilized with 100% methanol at −20 °C for two minutes.

CSK treatments were performed after washing cells with PBS (Thermo Fisher, 14190169) using 500 ul CSK buffer (10 mM PIPES, 100 mM NaCl, 3 mM MgCl2, 300 mM sucrose, 0.1% Triton X-100) with or without 300 ng/ml RNase A (Thermo Fisher, 12091021) at room temperature for 5 minutes. Cells were then fixed in 3% paraformaldehyde and washed twice in PBS. After fixation, coverslips were blocked in 2% BSA in PBS-T (0.1% Triton X-100 in PBS) for 1 hour and then incubated in primary antibodies against: TOP2A diluted 1:200 (Abcam, ab52934), fibrillarin diluted 1:150 (Santa Cruz Biotechnology, sc-166001), MYC diluted 1:200 (Abcam, ab32072), or TOP2B diluted 1:200 (Sigma, HPA024120) in PBS-T with 2% BSA. Coverslips were then washed three times in PBS-T and incubated in anti-rabbit (Abcam, ab205718) or anti-mouse (Abcam, ab205719) fluorescently conjugated secondary antibodies diluted 1:500 and DAPI diluted 1:1000 (Thermo Fisher, D1306) in PBS-T with 2% BSA. Finally, coverslips were washed three times in PBS-T and mounted using ProLong Diamond Antifade Mountant with DAPI (Thermo Fisher P36966) and sealed with nail polish or mounted using ProLong Diamond Antifade Mountant (Thermo Fisher P36970) and left to cure overnight.

TOP2A nucleolar to nucleoplasmic intensities were calculated using CellProfiler. Fibrillarin staining was used to identify nucleoli, and DAPI staining was used to identify the whole nucleus. Nuclei were masked with nucleolar objects to obtain nucleoplasmic signal. TOP2A intensity was measured in the nucleoli and nucleoplasm, and mean nucleolus intensity was divided by mean nucleoplasmic intensity for each cell.

Cells were imaged on Zeiss LSM 700, 710, 880, 980-Airy confocal microscopes. Graphs and statistical analysis were done on GraphPad Prism version 10.4.1.

### Fluorescence recovery after photobleaching (FRAP)

Recombinant SNAP-tagged TOP2A was labeled with SNAP-surface Alexa Fluor 488 as described above before being aliquoted and stored at –80 °C. 500 nM TOP2A, 10 ng/ul pUC19 plasmid DNA, and 3.5 μM MYC^WT^ were combined in 384-well black non-binding μCLEAR microplates (Grenier, 781906), and droplets were allowed to form and settle for 10 minutes before bleaching. Whole droplets were bleached with a 488 nm laser at 100% intensity for 75 iterations. Bleached droplet intensity was measured over 8 minutes, and droplet intensity was normalized to that of a second, unbleached droplet. Intensity measurements were taken using ImageJ. FRAP was performed on Zeiss LSM 980-Airy confocal microscope. Graphs and statistical analysis were done on GraphPad Prism version 10.4.1.

### Cover glass functionalization, sample preparation and fluorescent wide-field imaging

DNA-protein complexes were deposited on functionalized microscope coverslips following published protocols^43,73,74^. Coverslips and cover glasses were washed by sonication in a 2% Hellmanex solution, rinsed 3 times with MilliQ (MQ) water, dried with N_2_-gas, rinsed with acetone, submerged in Allyltrimethoxysilane (ATMS, 95%, Sigma-Aldrich), (3-aminopropyl) triethoxysilane (APTES,≥98%,Sigma-Aldrich) and Acetone (Sigma-Aldrich) at a 1:1:100 ratio for 2 hours. Prior to loading a sample, the coverslips were rinsed with MQ water and dried with N_2_-gas. An 8 μL sample was used per coverslip. A reaction consisting of TOP2A, MYC, and DNA in 50 mM Tris pH 8, 150 mM NaCl, 10 mM MgCl_2_, 0.5 mM DTT and 30 μg/mL BSA was setup by equilibrating buffer and protein on ice for 10 minutes. After adding the DNA, the reaction was incubated at 37 °C for 10 minutes and Sytox Orange (SYTOX™ Orange Nucleic Acid Stain, Invitrogen, S11368) was added. The sample was imaged immediately.

Images were taken with an inverted fluorescence microscope (Zeiss AxioObserver.Z1) equipped with a 63x oil immersion objective (NA=1.46, Zeiss), 1.6×optovar magnification changer, an iXon EMCCD camera (Andor) and an LDI-7 Laser Diode Illuminator (89 NORTH) was used. The sample was alternately excited with 640 nm, 555 nm or 470 nm light and filtered with Zeiss Filter set 50, 43 or 44, respectively.

Images were processed with a custom Python script. Briefly, the images were flat-field corrected (FFC), filtered with a gaussian and a top-hat filter and segmented by joining the result from a local thresholding algorithm and a difference-of-gaussian blob detector. The background was defined as the inverse of the puncta mask. Puncta statistics was extracted from the FFC images.

### Glycerol gradients and Western blotting

K562^MYC-AID^ cells were treated with 0.1 mM auxinole (MedChemexpress) for 24 hours, to block MYC degradation, or 500 μM auxin (3-indoleacetic acid, Sigma, I3750) for 90 minutes. Cells were counted using Countess 3 cell counter (Invitrogen) and 8 million cells were harvested, spun down, washed once in PBS (Thermo Fisher, 14190169), and washed once in HWB (20 mM HEPES pH 7.5, 137 mM NaCl). Cells were then lysed in HLB (10 mM HEPES pH 7.5, protease inhibitor tablet (cOmplete, EDTA-free protease inhibitor cocktail, Merck, 4693132001), 0.5 mM AEBSF (4-(2-Aminoethyl) benzenesulfonyl fluoride hydrochloride, Merck, A8456), 10 μM Leupeptin (Leupeptin trifluoroacetate salt, Merck, L2023), 1 μM Pepstatin A (Merck, P5318), and 7 mM NaF and passed through a 20 G needle 10 times to disrupt the cell membrane before being spun down for 5 minutes at 800 g. The resulting pellet was washed again with HLB before being spun again and resuspended in 0.2% sarkosyl, 20 mM HEPES pH 8, 133 mM NaCl, 2 mM MgCl_2_, 0.4 mM EDTA. Chromatin was sheared through passage with 25 G needle 7 times, sonication for 5 minutes, and treated for 20 minutes with 250 units benzonase (Merck, E8263) before being centrifuged to remove the remaining insoluble fraction. Samples were loaded into a 10-30% glycerol gradient (10 or 30% glycerol, 20 mM HEPES pH 8, 133 mM NaCl, 2 mM MgCl_2_, 0.4 mM EDTA) created in Polyclear Open Top Ultracentrifuge Tubes (Biocomp Instruments, 151-514A) using a BioComp gradient master, and spun for 16 hours at 35000 RPM at 4 °C in a Beckman Coulter Optima L-90K ultracentrifuge fitted with a SW41 rotor. Gradients were manually fractionated into 24 fractions and precipitated using 0.18% DOC for 10 minutes followed by the addition of 9% TCA for 30 minutes. The fractions were centrifuged at 21130 RCF for 10 minutes, washed twice in 25% acetone, and the protein pellet was resuspended in 1x Bolt LDS (Invitrogen, B007) with 5% β-mercaptoethanol.

Samples were loaded onto Bolt Bis-Tris Plus Mini Protein Gels (Invitrogen, NW04127BOX), ran for 70 minutes at 160 V, and transferred onto nitrocellulose membrane overnight. The membrane was blocked in 3% BSA in TBS-T. Incubation with anti-TOP2A (Abcam, ab52934) and anti-MYC (Abcam, ab32072) primary antibodies was done in 3% BSA in TBS-T diluted 1:10,000. Incubation with secondary antibodies (anti-rabbit: Abcam, ab205718 or anti-mouse Abcam, ab205719) was done in 3% BSA in TBS-T diluted 1:35000. The blot was imaged using SuperSignal West Femto Maximum Sensitivity Substrate ECL (Thermo Scientific, 34095) in a Biorad Chemidoc MP Imaging System. For the western blots shown in Extended Data Fig. 6, we used anti-beta Actin (Merck, A5441) primary antibody in 3% BSA in TBS-T at 1:10,000 concentration. Quantification was done using Image Lab Version 6.1.0 build 7.

### Coimmunoprecipitation of recombinant proteins

Coimmunoprecipitations (coIPs) with recombinant proteins were carried out by mixing 200 ng of full length TOP2A or C-terminally truncated TOP2A (ΔCTD) with 100 ng of MYC protein in PDB buffer (10 mM Tris-HCl pH 7.5, 100 M KCl, 0.5% NP40, 7.5% Glycerol, 200 μM EDTA, 250 μg/ml BSA, supplemented with protease and phosphatase inhibitors) on ice for 30 minutes. 100 ng of anti-TOP2A/B (Abcam, ab109524), anti-MYC (Abcam, ab32072) or IgG control (Santa Cruz, sc-2025) was added to the mixture followed by incubation on ice for 30 minutes. For each sample, 6 μL Protein A/G beads (Thermo Fisher, 88803) were blocked in PDB buffer with 5% skim milk powder for 1 hour at 4 °C, washed in PDB and added to the protein solution in 100 μL PDB followed by incubation for 1 hour at 4 °C. Washes were performed twice with each of the following buffers: PDB buffer with 5% skim milk powder, high-salt PDB buffer (250 mM KCl), and PDB buffer with 0.2% Sarkosyl and 5% skim milk powder. Final washes were performed once with 500 μL PDB buffer and once with 40 μL PDB. Proteins were eluted by incubating the beads at room temperature for 10 minutes in 15 μL of 1× Bolt LDS loading buffer (Thermo Fisher, B0007). After elution from beads, β-mercaptoethanol was added to a final concentration of 5%, and samples were heated at 70 °C for 10 minutes. Protein samples were examined by western blotting. Briefly, samples were run on 4%–12% Bis-Tris Bolt protein gels (Thermo Fisher, NW04122BOX) for 1 hour, transferred overnight to nitrocellulose membranes, probed for 6 hours with antibodies against TOP2A and MYC proteins, and imaged by chemiluminescence using ECL chemiluminescence reaction (Thermo Fisher, 34075). All densitometric quantifications of immunoblots were carried out using Image Lab software from Bio-Rad.

### Coimmunoprecipitation of nuclear extract

HCT116^TOP1-AID^ cells were seeded and allowed to grow for two days before being treated with 500 μM auxin for 90 minutes or treated with auxinole 24 hours before harvesting. Protein A/G beads (Thermo Fisher, 88803) were washed in RIPA-IP 137 (137 mM NaCl, 50 mM Tris-HCl pH 7.5, 10% NP40) and incubated in RIPA-IP 137 + 1% BSA for 30 minutes at 4 °C. Beads were then washed again and incubated in 1 μg of anti-TOP1 primary antibody (Abcam, ab109374) and anti-Flag primary antibody (Sigma, F1804) or anti-TOP2A primary antibody (Abcam, ab52934) for 3 hours at 4 °C. Nuclear extracts from 2 million cells were harvested as described above and incubated with anti-TOP1 and anti-FLAG bound to beads to remove residual TOP1 for 3 hours at 4 °C. Supernatant from the resulting IP was then incubated overnight with anti-TOP2A bound beads at 4 °C. Beads were washed once with RIPA-IP 137 and twice with RIPA-IP 250 (250 mM NaCl, 50 mM Tris-HCl pH 7.5, 10% NP40). Proteins were eluted from beads at room temperature in 1x Bolt LDS loading buffer (Thermo Fisher, B0007). After elution from beads, β-mercaptoethanol was added to a final concentration of 5%, and samples were heated at 70 °C for 10 minutes. Samples were probed using Western blotting as described above.

### Single molecule tracking (SMT)

Halo-TOP2A HCT116^MYC-AID^ cells were plated in 4-well Lab-Tek II Chamber slides (Nunc, 155382) in phenol-free high-glucose DMEM media (Thermo, 31053028) with 10% FBS and GlutaMAX to achieve 40-50% confluence upon imaging 48 hours later. To image nucleoli, we transfected with a 1:99 ratio of GFP-Nucleolin vector (Addgene, 28176) and an empty vector (pCDNA4/TO, Thermo, V102020) 24 hours after plating using Lipofectamine 3000 (Thermo, L3000008), according to the manufacturer’s protocol. The TOP2A was then labeled with 100 nM photo-activatible Janelia Fluor (JF) 549 Halo ligand and 10 nM JF-646 Halo ligand for 30 minutes at 37 °C in the cell culture incubator. The cells were then washed three times with warmed PBS and incubated for 15 minutes in growth media. This washing process was repeated once before the media was replaced again with growth media to remove all excess Halo ligand. If cells were treated with 500 μΜ auxin, the auxin was included during both 15 minute incubations and in the final media suspension to ensure knockdown had occurred for at least 60 minutes before imaging.

Cells were imaged using a custom-built microscope able to perform single molecule imaging by HILO illumination as previously described^28^. Nuclear TOP2A distribution was determined from the JF-646 channel, enabling distinction between the nucleolus and the nucleoplasm based on the staining intensity. Photo-activation of the JF-549 ligand was achieved by tuning the power of a 405 nm laser to ensure 2-5 molecules of JF-549 ligand were photo-active in each frame. For each video, 3000 frames were captured at 100 frames per second. Measurements for the nuclear, nucleolar and nucleoplasmic regions were determined by only analyzing the region of interest after manually segmenting based on the TOP2A JF646 signal. The path of individual molecules from the SMT video was determined using the ImageJ Trackmate plugin^75^ without gaps being allowed and with 1 μm maximum linking distance. Tracks were classified into fast, slow and bound molecules using the vbSPT algorithm^30^ excluding tracks shorter than 4 frames.

To calculate diffusional anisotropy, we removed bound track segments and calculated the angle between consecutive displacements. The anisotropy index was calculated as the ratio of the number of backward jumps (with an angle between jumps in the [150°, 210°] range) over the number of forward jumps (angles in the range [-30°, 30°]). Plotting the anisotropy index as a function of the distance run by the molecule allows discrimination between exploration modes. Error bars were calculated as standard deviation from 100 random subsampling composed by 50% of the original data. Analysis of diffusion coefficients and fractions at the single cell level was performed by analyzing the distribution of displacements with a three-component diffusion model, as described previously^28^.

### Quadruple-trap (Q-TRAP) optical tweezers

Single molecule TOP2A decatenation assays were performed using the C-TRAP optical tweezers (Lumicks). Two molecules of lambda DNA with biotinylated ends (brand) were each bound to two SPHERO streptavidin-coated polystyrene particles (Spherotech, SVP-40-5) held by optical tweezers. These particles were manipulated to intertwine the two DNA molecules forming five DNA crossovers. This structure was then moved into a channel that had been passivated overnight in passivation buffer (50 mM Tris pH 8, 150 mM NaCl, 10 mM MgCl_2_, 0.5% pluronic, 0.1% BSA). This channel contained 10-50 nM SNAP-TOP2A labeled with SNAP-647 ligand, 100-300 nM MYC or equivalent BSA, 50 mM Tris pH 8, 150 mM NaCl, 10 mM MgCl_2_, 450 nM BSA, 0.5 mM DTT, 50 μΜ Sytox Orange, and 2 mM ATP where required.

The exact time of entrance of the DNA crossover into the protein channel was determined by the background photon count in the TOP2A channel. The exact time of decatenation was determined by the change in the force acting on the polystyrene particles. Droplet formation at the crossover was measured by comparing the intensity of TOP2A at the DNA crossover, subtracting the background TOP2A signal surrounding the crossover and comparing this value to a threshold for TOP2A crossover binding.

### Stimulated emission depletion (STED) microscopy

HCT116 cells were plated onto 18 mm coverslips (Marienfeld, 630-2200) in 12-well plates, and grown to 50-60% confluency. After the indicated treatment, cells grown on glass coverslips (Marienfeld, 630-2200) were fixed in 3% paraformaldehyde (PFA, Pierce, 28908) in PBS for 5 minutes at room temperature. After washing with PBS, cells were permeabilized in methanol for 3 minutes at −20 °C, before washing again in PBS and blocking in 2% BSA (Sigma, A7906) in PBS for 30 minutes at room temperature. Proteins were fluorescently labeled by sequential incubation with primary (1:200 dilution) and fluorescently labeled secondary (1:40 0 dilution) antibodies for 1 hour each in 0.5% BSA/PBS. After washing, coverslips were mounted onto slides in a hard-setting ProLong Diamond Antifade Mountant (Thermo, P36970). Multi-color STED imaging was performed on a Leica SP8X system, equipped with a tunable white-light laser (excitation 470-670 nm), high-power green/orange/red depletion lasers (592, 660, 775 nm), a chromatically optimized 100x/1.4 STED WHITE oil immersion objective, and single photon sensitive detectors (HyD-SMDs). Nuclear protein structures were imaged sequentially frame by frame (1024×1024 pixels) at a speed of 200 lines per second (4-line averages) with a pixel size of 25 nm and pinhole settings of 0.9-1.0 Airy units. STED images were deconvoluted using the Huygens 24.10 software (Scientific Volume Imaging) before data analysis. Images were analyzed using custom Matlab scripts. Briefly, the central points of nucleoplasmic puncta were determined at a sub-pixel resolution. Then, the signal from the observed channel surrounding the detected puncta is averaged and plotted.

### Quantitative PCR (q-PCR)

HCT116 cells were treated as indicated, then RNA was purified using NucleoSpin RNA columns (Macherey-Nagel, 740955) according to kit instructions. Equal amounts of RNA resuspended in 13 ul H_2_O were mixed with 1 ul of 10 mM dNTP (10 mM each of dATP, dCTP, dGTP, dTTP; Promega, U1205, U1215, U1225, U1235) and 0.5 ul of random hexamers (Promega, C1181). This mixture was heated at 65 °C for 5 minutes, then put on ice for 1 minute. 4 ul RT buffer, 0.3 ul SuperScript IV Reverse Transcriptase (both Thermo, 18090200), 1 ul 0.1 M DTT (Sigma, 43815) and 0.2 ul of RNAsin Ribonuclease inhibitor (Promega, N2515) were added. The solution was incubated at room temperature for 10 minutes, followed by 50 °C for 30 minutes to transcribe cDNA, and at 80 °C for 10 minutes to stop the reaction. The cDNA was diluted and quantitative PCR was performed using Fast SYBR Green Master Mix (Thermo, 4385612) with primers for the internal transcribed sequence (ITS) of the 47S pre-ribosomal RNA (F: CCG TGG CCT TAG CTG CTC GC, R: CCC ACT TAA CTA TCT TGG GCT G) and beta-2-microglobulin (B2M; F: GAA ACC TTC CGA CCC CTC T, R: GCC AGA CGA GAC AGC AAA C). The ΔΔCt method was used to measure ITS expression relative to B2M normalized to the control condition.

### EU labeling of nascent RNA

K562^MYC-AID^ cells were seeded and allowed to grow for 48 hours before being treated with 500 μM auxin (3-indoleacetic acid, Sigma, I3750) for 90 minutes or auxinole 24 hours before harvesting. Cells were additionally incubated with 5-ethynyl uridine (EU) (from Click-iT™ RNA Alexa Fluor™ 488 Imaging Kit, Thermo, C10329) for an hour without removing auxin/auxinole. Cells were then washed in PBS once before being fixed in 2% paraformaldehyde (PFA, Pierce, 28908) in PBS for 10 minutes at 4 °C before being washed in PBS. Cells were resuspended in provided Click-it Reaction buffer and incubated for 30 minutes, washed in reaction rinse buffer and resuspended in 0.1% Tween-20 in PBS. EU incorporation was measured on an ID7000™ Spectral Cell Analyzer. EU high or low cells were determined using a cutoff that excluded >99% of cells measured using a control cell population not treated with EU. Analysis and histogram were generated using FlowJo version 10.8.1 and statistical analysis was performed on GraphPad Prism version 10.4.1.

### TOP2 covalent adduct detection coupled to sequencing (TOP2 CAD-seq)

3×10^7^ (HCT116^MYC-AID^) cells were treated (in biological triplicates) with auxin followed by 10 μM MG132 for 30 minutes and 100 μM Etoposide in the last 6 minutes to trap TOP2cc. Cells were lysed in 4.5 mL of M buffer (9.3 mM Tris-HCl pH 6.5, 18.6 mM EDTA, 5.59 M guanidine thiocyanate, 0.93% DTT, 0.93% Sarcosyl, 3.72% Triton X-100) and briefly sonicated with Bandelin probe sonicator at 20% amplitude for 1 second. DNA covalent adducts were precipitated with 50% EtOH at −20 °C, centrifuged at 14000 rpm at 4 °C and pellets were washed thrice in wash buffer (20 mM Tris-HCl pH 7.5, 50 mM NaCl, 1 mM EDTA, 50% EtOH). Ethanol leftovers were aspirated and pellets were dried for 5 minutes and resuspended in 1 ml of EB-SDS 0.05% (10 mM Tris-HCl pH 8, 0.05% SDS). After 30 minutes incubation by gentle agitation, 500 μl of each sample were further fragmented with benzonase (0.02 U/μg) at 37 °C for 15 minutes. The reaction was stopped with 50 mM EDTA and 0.1% SDS. For the immunoprecipitation 3.5 μg of 1:1 mixture of anti-TOP2A (sc166934 and ab52934) antibodies were mixed with 25 μl Protein A/G magnetic beads (Pierce, 88803) and incubated at 4 °C for 4 hours with rotation. Beads were washed once with ice-cold PBS and DNA covalent adducts from 1.5 × 10^7^ cells were added to the Protein A/G-antibody complexes and incubated overnight at 4 °C with rotation. Samples were washed once with RIPA buffer, once with RIPA buffer containing 300 mM NaCl, once with LiCl-SDS 0.1% buffer (10 mM Tris-HCl pH 8, 1 mM EDTA pH 8, 250 mM LiCl, 0.5% NP40, 0.5% Na-Deoxycholate, 0.1% SDS) and twice with TE-SDS 0.1% (10 mM Tris-HCl pH 8, 1 mM EDTA pH 8, 0.1% SDS). The beads were then resuspended in 100 μl TE plus 0.5% SDS supplemented with proteinase K (500 μg/ml) and incubated for 4 hours at 60 °C. Samples were then purified by QIAquick PCR purification Kit.

### Library preparation and sequencing of TOP2 CAD-seq

DNA from TOP2A CAD samples was quantified with the Qubit dsDNA HS Assay Kit (Thermo Fisher, Q33230). To cleave off covalently bound tyrosyl residues from TOP2A, the samples were additionally treated with ExoVII (NEB, M0379S) (0.5 U per 10 ng of DNA) and purified by PCR purification Kit (QIAGEN, 28106). Sequencing libraries were created according to the ThruPLEX DNA-seq kit protocol (Takara, R400676). Size selection was performed in the range of 200– 700 bp with AMPure XP beads (Beckman, A63880) and confirmed using the Agilent High Sensitivity DNA Kit (Agilent, 5067–4626) on the Agilent 2100 Bioanalyzer. Libraries were pooled and sequenced using the NextSeq 1000/2000 P2 XLEAP-SBS Reagent Kit (100 Cycles) (Illumina, 20100987). The sequencing run was Pair End and Dual Index with 2 x 50 bp reads.

### Data analysis

The generated fastq files were quality controlled with FastQC (https://www.bioinformatics.babraham.ac.uk/projects/fastqc/) and MultiQC^76^, trimmed with cutadapt (http://journal.embnet.org/index.php/embnetjournal/article/view/200/479), aligned to hg38 reference genome with bowtie2^77^, deduplicated, sorted and indexed using Samtools^78^ and Picard (http://broadinstitute.github.io/picard). RPM normalized BigWig files were generated with bamCoverage^79^. The coverage matrices, profiles and heatmaps of short reads’ average distribution near TSSs and along normalized gene bodies were generated by deeptools commands computeMatrix and plotHeatmap. Only the protein coding genes from the Ensembl #76 database^80^ were used for generating profiles and heatmaps. The expression of these genes was determined based on previously published RNA-seq in HCT116 cells^12^. Expressed (FPKM > 0) genes have been ranked in descending order and top 1500 selected for Fig. 5h. Peaks were called using macs3 callpeak and peak summits were used for plotting.

### Quantification and statistical analysis

All image analysis for nuclear localization of TOP2A, γH2AX, and droplet imaging experiments (for 2 μM and 0.5 μM concentrations) were done using CellProfiler version 4.2.8 and statistical analysis was performed in GraphPad Prism version 10.4.1. Western blot analysis was performed using Image Lab Version 6.1.0 build 7, and plots and statistical analysis was performed on GraphPad Prism.

## Acknowledgements

This work was supported by the ERC Consolidator Grant (project no. 101088643 to L.B.), Knut och Alice Wallenbergs Stiftelse (KAW 2022.0380 and KAW 2022.0189 to L.B.), the Swedish Research Council (2021-02630 to L.B.), Cancerfonden (21 1771 Pj 01 H to L.B.), and KI Consolidator (2-190/2022 to L.B.). This work was supported by ERC Consolidator Grant (project no. 866238) and the Swedish Research Council (2020–03400) to FW. E.P. acknowledges funding from the Area of Advance nano at Chalmers. D.M. acknowledges funding from Worldwide Cancer Research (Grant Reference number 22-0116) and from AIRC under the IG 2018-ID:21897, IG 2023-ID:28792. V.L. acknowledges the support of the Integrated Structural Biology platform of the Strasbourg and Instruct-ERIC center IGBMC-CBI supported by FRISBI (ANR-10-INBS-0005), and funding from the Interdisciplinary Thematic Institute IMCBio+, supported by IdEx Unistra (ANR-10-IDEX-0002), and by SFRI-STRAT’US project (ANR-20-SFRI-0012) under the framework of the France 2030 Program. B.L.D. is a doctoral fellow supported by EUR IMCBio (ANR-17-EURE-0023). D.C. was supported by the Helge Ax:son Johnson Foundation (F22-0197), the Erik and Edith Fernström Foundation for Medical Research (2022-00603) and the UTokyo-KI LINK programme.

We thank the SciLifeLab/NMI Advanced Light Microscopy unit for STED support. The computations and data storage were enabled by resources in project [SNIC 2018/8-390], provided by the Swedish National Infrastructure for Computing (SNIC) at UPPMAX, partially funded by the Swedish Research Council through grant agreement no. 2018-05973. Part of this work was facilitated by the Protein Science Facility at Karolinska Institutet/SciLifeLab (https://ki.se/psf) and the Karolinska Genome Engineering Facility (KGE). Cell imaging was performed at the Biomedicum Imaging Core (BIC) with support from Karolinska Institutet.

We thank Dr. Johannes Zuber for sharing reagents and helpful advice. We thank Dr. Camilla Björkegren, Dr. Kristian Jeppsson, Dr. Arne Lindqvist, Dr. Fedor Kouzine, Dr. Keir Neuman, Dr. Katsuhiko Shirahige, and Dr. David Levens for helpful advice and critical discussion. We thank Dr. Katsunori Fujiki, Dr. Florian Salomons, Dr. Hans Blom, Dr. Ola Larsson, Dr. Solenne Bleuse and Dr. Claudia Kutter for sharing instruments and technical expertise. The photo-activatable JF549 Halo-tag was generously gifted by Dr. Luke Lavis.

## Illustrations

Illustrations in Fig. 2a, Fig. 4a,d,g, Fig. 5c,g, Fig. 6 and Extended Data Fig. 5d were created with Biorender.com.

## Data availability

The data for this study have been deposited in the Gene Expression Omnibus with accession number GSE298118, and will be available upon publication. All additional information and data reported in this paper are available from the corresponding author upon request.

## Declaration of generative AI and AI-assisted technologies in the writing process

During the preparation of this work the authors used ChatGPT o1 in order to improve the readability of the manuscript. After using this tool/service, the authors reviewed and edited the content as needed and take full responsibility for the content of the published article.

## Author contributions

Conceptualization: D.P.C., K.J., D.M., L.B. Methodology: D.P.C., K.J., A.L., C.M., V.K., M.M., F.W., D.M., L.B. Software: D.P.C., K.J., C.M., V.K., D.M. Formal Analysis: D.P.C., K.J., C.M., V.K. D.M. Investigation: D.P.C., K.J., A.L., C.M., V.K., M.M., E.I., E.P., B.J., B.S.L.D. Resources: V.L., F.W., D.M., L.B. Writing - Original Draft: D.P.C., K.J., L.B. Writing - Review and Editing: D.P.C., K.J., A.L., C.M., V.K., M.M., E.I., E.P., B.J., B.S.L.D., V.L., F.W., D.M., L.B. Visualization: D.P.C., K.J., C.M., V.K., L.B. Supervision: D.P.C., V.L., F.W., D.M., L.B. Funding acquisition: D.P.C., V.L., F.W., D.M., L.B.

---

**Extended Data Fig. 1.**
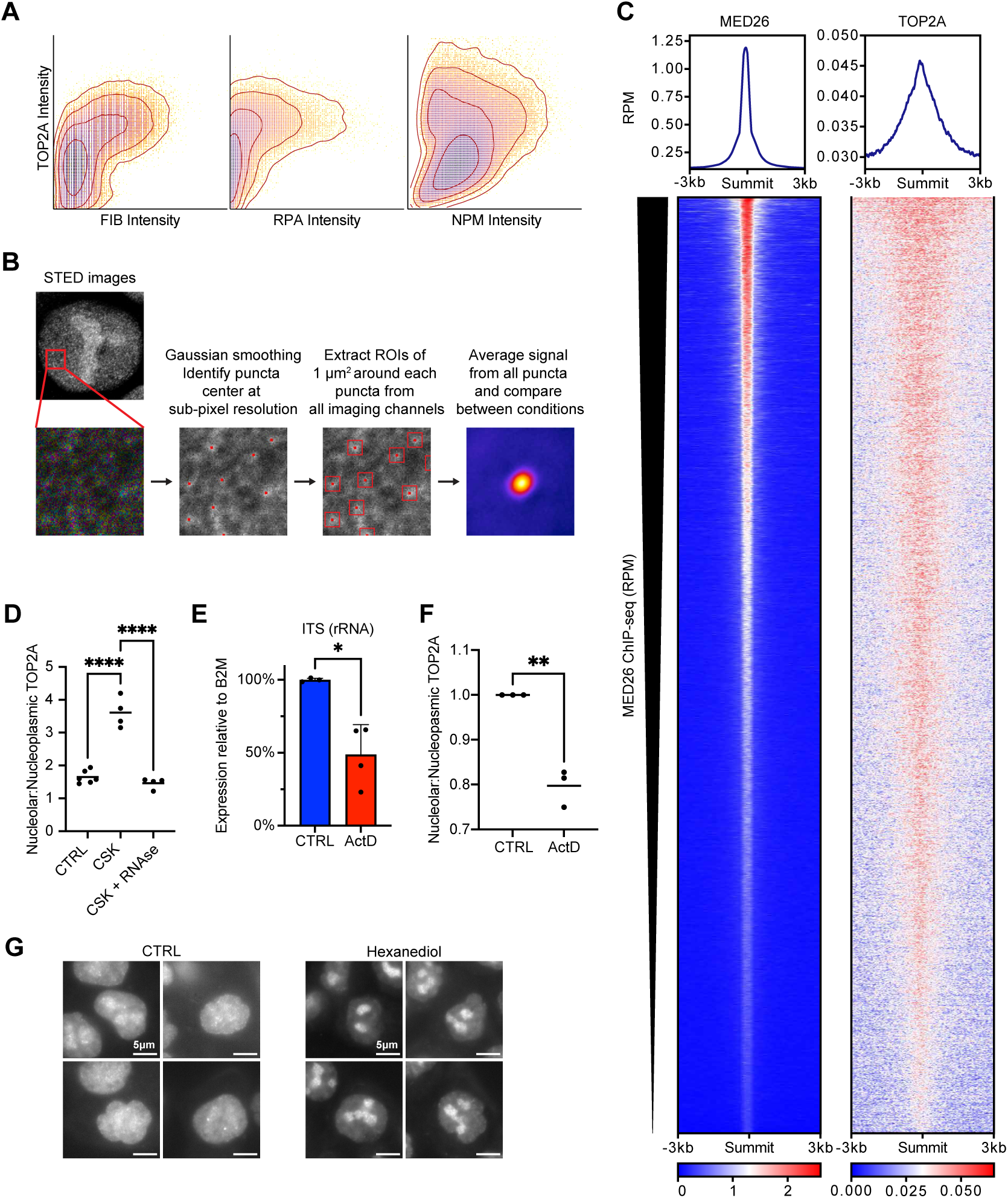
TOP2A nuclear association and dynamics. **a**, Correlation analysis showing pixel intensity for the protein pairs from images in Fig. 1a. Left, FIB *vs* TOP2A. Middle, RPA194 *vs* TOP2A. Right, NPM *vs* TOP2A. **b**, Scheme of method for puncta detection. **c**, Heatmaps of TOP2A and MED26 ChIP-seq at MED26 peak summits +/− 3 kb. Data retrieved from publicly available datasets^5,19^. **d**, Quantitation of nucleolar:nucleoplasmic ratio of TOP2A signal normalized to CTRL after treatment with CSK buffer to remove soluble proteins +/− RNAse. Note that the increased ratio after CSK is due to low fluorescent signal. Average of four independent experiments. 550 CTRL, 423 CSK-treated, and 306 CSK+RNAse-treated cells were measured. ****p < 0.0001 (one-way ANOVA, Tukey correction). **e**, RNA expression of 47S pre-rRNA precursor transcript measured by qPCR after 60 minutes treatment with 5 nM ActD, normalized to Beta-2-microglobulin transcript expression. *p < 0.05 (Paired t-test). **f**, Quantitation of nucleolar:nucleoplasmic ratio of TOP2A signal normalized to CTRL after 1 hour of 5 nM ActD treatment of HCT116 cells. Three independent experiments were performed. 571 CTRL cells and 512 ActD-treated cells were measured. **p < 0.001 (unpaired t-test). **g**, Representative images of Halo-TOP2A HCT116 cells with fluorescently labelled TOP2A after 5-15 minutes treatment with 4% 1,6-hexanediol. Scale bar = 5 μm.

**Extended Data Fig. 2.**
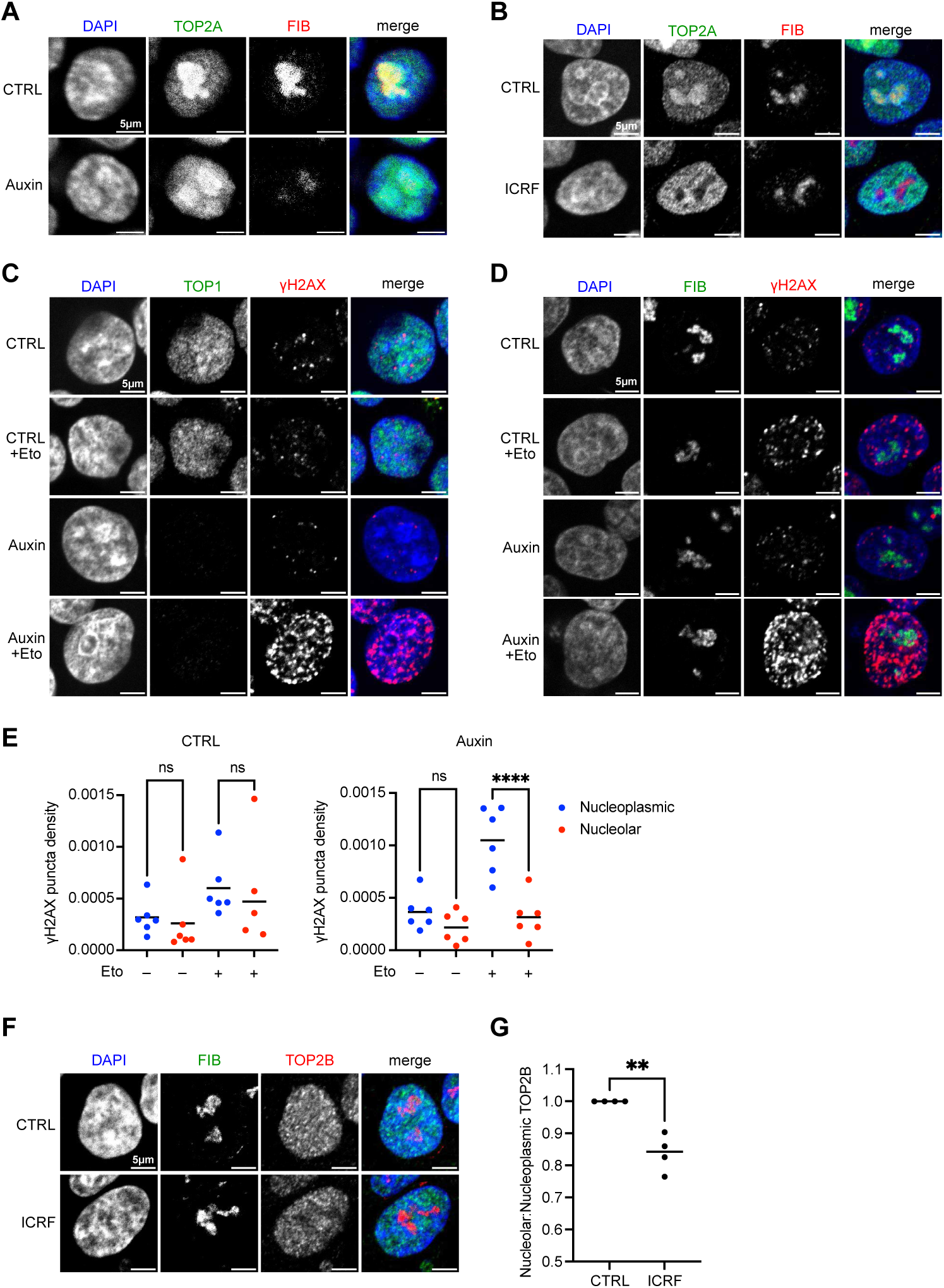
TOP2-dependent DNA damage is enriched in the nucleoplasm. **a**, Representative confocal images of HCT116^TOP1-AID^ cells +/− auxin treatment immunostained for TOP2A and FIB, and DNA labelled with DAPI. Scale bar = 5 μm. Quantitated in Fig. 1e. **b**, Representative confocal images of HCT116 cells +/− ICRF-193 treatment immunostained for TOP2A and FIB, and DNA labelled with DAPI. Scale bar = 5 μm. Quantitated in Fig. 1e. **c**,**d**, Representative images of HCT116^TOP1-AID^ cells +/− auxin and +/− Eto treatment immunostained for γH2AX, TOP1 (A) or FIB (B) and DNA labelled with DAPI. Quantitated in Fig. 1f. **e**, Quantitation for experiment in **d**, where the γH2AX signal is separated into nucleolar and nucleoplasmic based on overlap with nucleolar marker FIB. 598 CTRL untreated cells, 576 CTRL Eto-treated cells, 566 auxin untreated cells, and 632 auxin Eto-treated cells were measured. ****p < 0.0001 (ordinary one-way ANOVA with Šidák correction). **f**, Representative images of HCT116 cells +/− ICRF-193 treatment immunostained for TOP2B and FIB, and DNA labelled with DAPI. **g**, Quantitation of nucleolar:nucleoplasmic ratio of TOP2B signal in HCT116 cells treated +/− ICRF. Four independent experiments were performed, normalized to respective CTRL conditions. 327 CTRL cells and 425 ICRF-treated cells were measured. **p < 0.01 (unpaired t-test).

**Extended Data Fig. 3.**
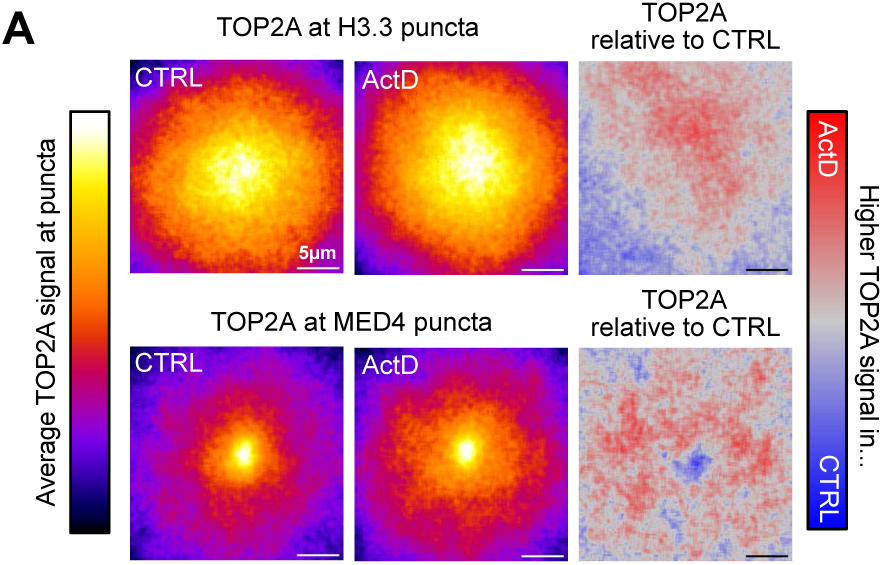
STED analysis of TOP2A overlapping puncta. **a**, Average TOP2A signal at H3.3 (top) and MED4 (bottom) puncta from STED images after control (left) or 5 nM Actinomycin D treatment for 1 hour (center), and difference between both conditions (right). Signal averaged from three independent experiments. 7273-14486 puncta per condition. Quantitated in Fig. 2g.

**Extended Data Fig. 4.**
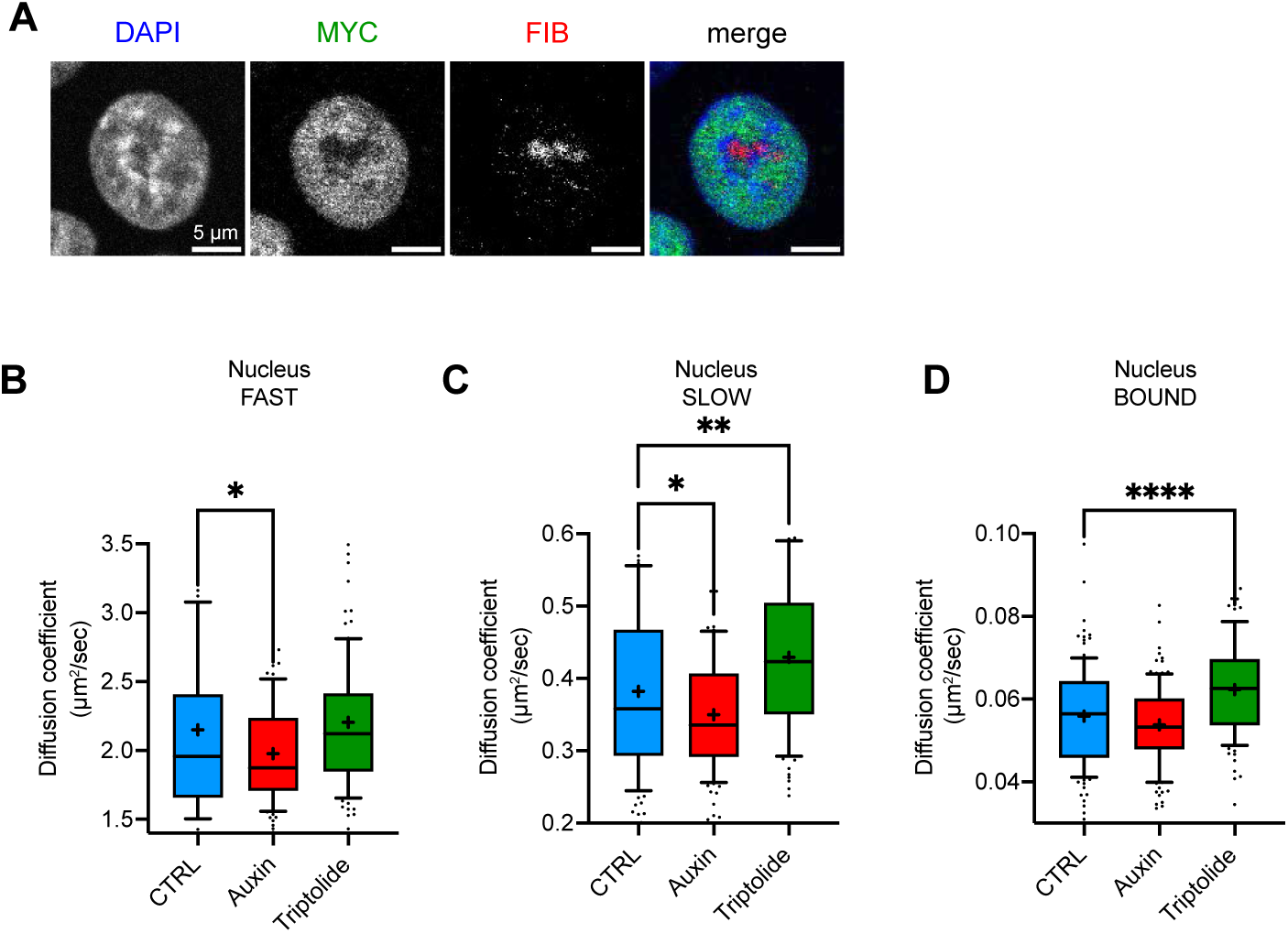
SMT analysis of TOP2A diffusion. **a**, Representative images of HCT116 cells immunostained for MYC and FIB, and DNA labelled with DAPI. **b**-**d**, Diffusion coefficients of Fast (**b**), Slow (**c**), and Bound (**d**) nuclear fractions from individual Halo-TOP2A HCT116^MYC-AID^ cells treated with control, auxin or triptolide (5 μM for 60 minutes). 110-129 cells for each condition from three independent experiments. ****p < 0.0001; **p < 0.01; *p < 0.05 (ANOVA, Dunnett’s multiple comparison test).

**Extended Data Fig. 5.**
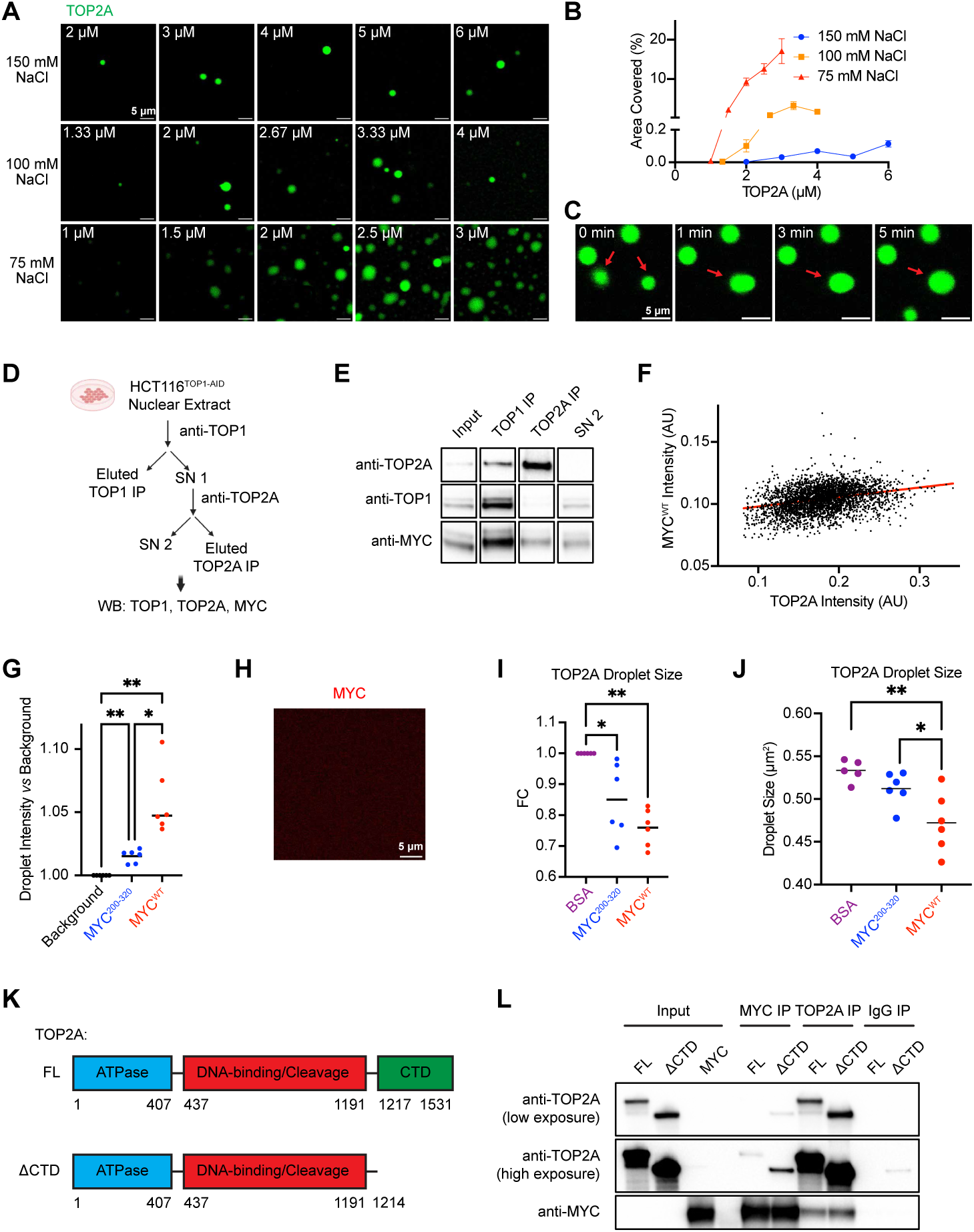
Characterization of TOP2A droplet +/− MYC *in vitro*. **a**, Representative images of SNAP-TOP2A droplets labelled with AF488 ligand at different protein and salt concentrations. Scale bar = 5 μm. **b**, Quantitation of percentage of area covered by SNAP-TOP2A droplets shown in Extended Data Fig. 5a. **c**, Example of TOP2A droplets fusing. **d**,**e**, HCT116^TOP1-AID^ cells are treated with auxin to degrade TOP1. Isolated nuclear extracts are incubated with anti-TOP1 (to remove traces of TOP1), the supernatant (SN) immunoprecipitated with anti-TOP2A, followed by western blot (WB) with anti-TOP1, -TOP2A and -MYC (**d**). Representative WB from two independent experiments is shown (**e**). **f**, Correlation between the intensity of SNAP-TOP2A and SNAP-MYC^WT^ in the same droplets. **g**, Intensity of droplets with SNAP-MYC^WT^ and SNAP-MYC^200–320^ *vs* background. **h**, Representative images of SNAP-MYC^WT^ labelled with AF647 showing that it does not form droplets under the conditions used in this study. Scale bar = 5 μm. **i**, Normalized quantitation of 2 μM SNAP-TOP2A droplet size +/− 3.5 μM BSA, MYC^WT^ or MYC^200–320^ in the presence of scDNA. Average of four independent experiments. **p < 0.01 (one way ANOVA, Dunnett’s test). **j**, Quantitation of 25 nM SNAP-TOP2A puncta area +/− 100 nM BSA, MYC^WT^ or MYC^200–320^ in presence of scDNA after deposition onto silanized slides. Average of 5-6 independent experiments. **p < 0.01; *p < 0.05 (ANOVA, Tukey’s multiple comparisons test). **k**, Schematic of full-length TOP2A (FL) and CTD-truncated TOP2A mutant (ΔCTD). **l**, Co-IP of recombinant full-length TOP2A or truncated TOP2A mutant with MYC using anti-TOP2A, anti-MYC or IgG, as negative control. Representative WB from three independent experiments.

**Extended Data Fig. 6.**
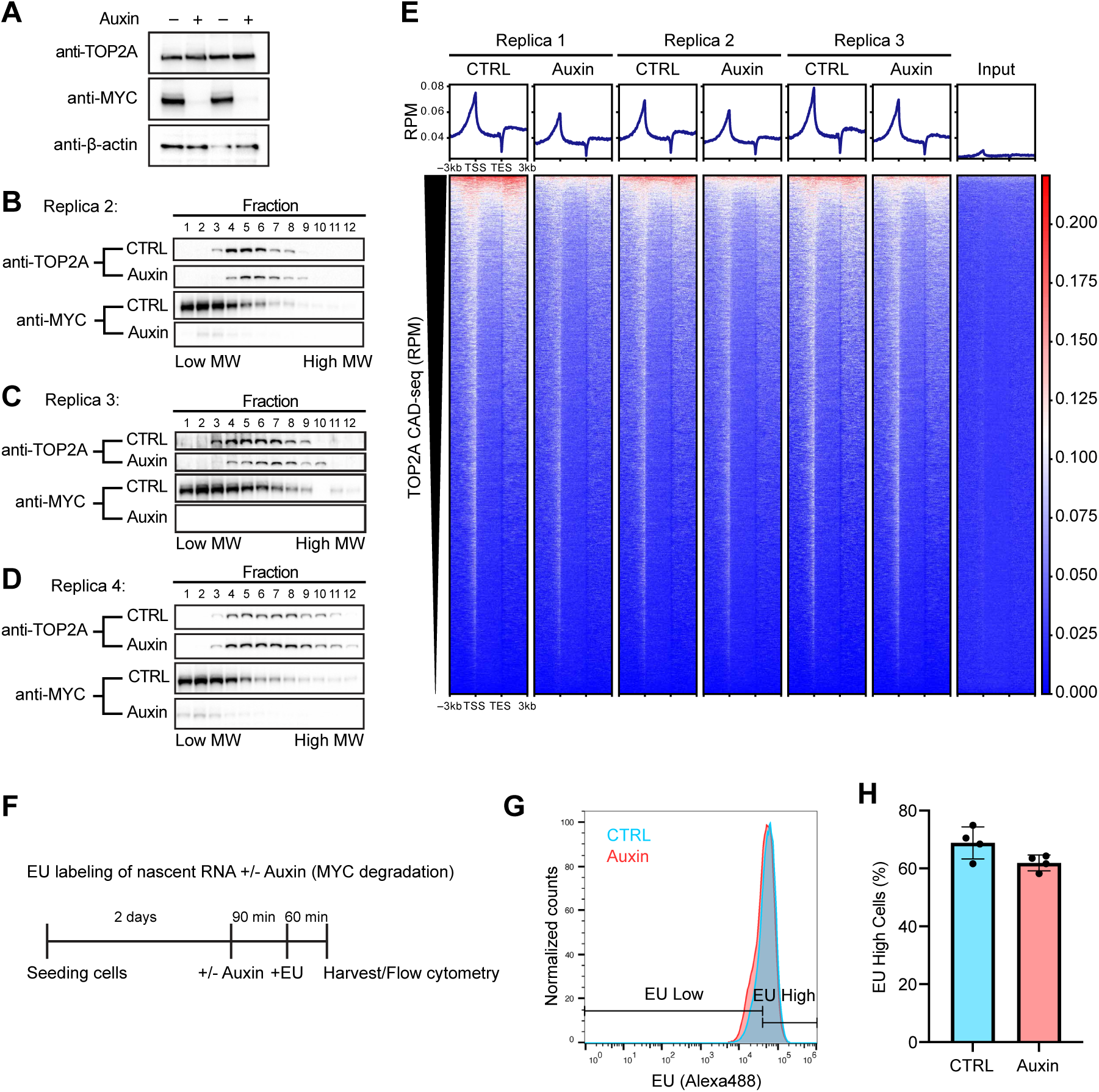
TOP2A complex size and activity in cells. **a**, Representative western blot of MYC and TOP2A from K562^MYC-AID^ cell lysates +/− 500 μΜ auxin treatment for 90 minutes prior to glycerol gradient demonstrating >95% knockdown of MYC protein. Two independent experiments were performed. **b**-**d**, Replicates of glycerol gradients from K562^MYC-AID^ cell lysates +/− 500 μM auxin treatment for 90 minutes. **e**, Heatmaps of TOP2A CAD-seq enrichment from three independent experiments across the gene bodies in HCT116^MYC-AID^ cells +/− auxin. Input DNA used as negative control. TSS refers to the transcription start site and TES refers to the transcription end site. Related to Fig. 5g,h. **f**, Scheme of EU experiment where K562^MYC-AID^ cells are pulsed with EU for 60 minutes to measure transcription levels upon MYC degradation by 90 minutes of auxin treatment. **g**, Representative histogram of EU incorporation by K562^MYC-AID^ cells. **h**, Quantitation of percent of EU high cells +/− auxin. Average of four independent experiments. ns: p > 0.05 (unpaired t test).

## References

1. Patange, S. et al. MYC amplifies gene expression through global changes in transcription factor dynamics. Cell Reports 38, 110292 (2022).

2. Liu, L. F. & Wang, J. C. Supercoiling of the DNA template during transcription. Proceedings of the National Academy of Sciences 84, 7024–7027 (1987).

3. Ma, J., Bai, L. & Wang, M. D. Transcription under torsion. Science (New York, N.Y.) 340, 1580–1583 (2013).

4. Pommier, Y., Sun, Y., Huang, S.-Y. N. & Nitiss, J. L. Roles of eukaryotic topoisomerases in transcription, replication and genomic stability. Nature reviews. Molecular cell biology 17, 703–721 (2016).

5. Das, S. K. et al. MYC assembles and stimulates topoisomerases 1 and 2 in a “topoisome.” Mol Cell 82, 140–158.e12 (2022).

6. Linka, R. M. et al. C-terminal regions of topoisomerase IIalpha and IIbeta determine isoform-specific functioning of the enzymes in vivo. Nucleic acids research 35, 3810–3822 (2007).

7. Lane, A. B., Giménez-Abián, J. F. & Clarke, D. J. A novel chromatin tether domain controls topoisomerase IIα dynamics and mitotic chromosome formation. The Journal of cell biology 203, 471–486 (2013).

8. Jeong, J., Lee, J. H., Carcamo, C. C., Parker, M. W. & Berger, J. M. DNA-Stimulated Liquid-Liquid phase separation by eukaryotic topoisomerase ii modulates catalytic function. Elife 11, e81786 (2022).

9. Shintomi, K. & Hirano, T. Guiding functions of the C-terminal domain of topoisomerase IIα advance mitotic chromosome assembly. Nat Commun 12, 2917 (2021).

10. Boija, A. et al. Transcription Factors Activate Genes through the Phase-Separation Capacity of Their Activation Domains. Cell 175, 1842–1855.e16 (2018).

11. Du, M. et al. Direct observation of a condensate effect on super-enhancer controlled gene bursting. Cell 187, 331–344.e17 (2024).

12. Baranello, L. et al. RNA Polymerase II Regulates Topoisomerase 1 Activity to Favor Efficient Transcription. Cell 165, 357–371 (2016).

13. Hell, S. W. & Wichmann, J. Breaking the diffraction resolution limit by stimulated emission: stimulated-emission-depletion fluorescence microscopy. Opt. Lett. 19, 780–2 (1994).

14. Stanek, D. et al. Non-isotopic mapping of ribosomal RNA synthesis and processing in the nucleolus. Chromosoma 110, 460–470 (2001).

15. Ray, S. et al. Topoisomerase IIα promotes activation of RNA polymerase I transcription by facilitating pre-initiation complex formation. Nature communications 4, 1598 (2013).

16. Britton, S., Coates, J. & Jackson, S. P. A new method for high-resolution imaging of Ku foci to decipher mechanisms of DNA double-strand break repair. J. Cell Biol. 202, 579–595 (2013).

17. Herrero-Ruiz, A. et al. Topoisomerase IIα represses transcription by enforcing promoter-proximal pausing. Cell Reports 35, 108977 (2021).

18. Cho, W.-K. et al. Mediator and RNA polymerase II clusters associate in transcription-dependent condensates. Science 361, 412–415 (2018).

19. Trzaskoma, P. et al. 3D chromatin architecture, BRD4, and Mediator have distinct roles in regulating genome-wide transcriptional bursting and gene network. Sci. Adv. 10, eadl4893 (2024).

20. Christensen, M. O. et al. Dynamics of human DNA topoisomerases IIalpha and IIbeta in living cells. The Journal of cell biology 157, 31–44 (2002).

21. Morotomi-Yano, K. & Yano, K. Nucleolar translocation of human DNA topoisomerase II by ATP depletion and its disruption by the RNA polymerase I inhibitor BMH-21. Sci. Rep. 11, 21533 (2021).

22. Itoh, Y. et al. 1,6-hexanediol rapidly immobilizes and condenses chromatin in living human cells. Life Sci Alliance 4, e202001005 (2021).

23. Boeynaems, S. et al. Phase Separation of C9orf72 Dipeptide Repeats Perturbs Stress Granule Dynamics. Mol. Cell 65, 1044–1055.e5 (2017).

24. Yamazaki, H., Takagi, M., Kosako, H., Hirano, T. & Yoshimura, S. H. Cell cycle-specific phase separation regulated by protein charge blockiness. Nat Cell Biol 24, 625–632 (2022).

25. O’Donnell, A. C. & Berger, J. M. Structural Mechanisms of Topoisomerase-Targeting Drugs. Annu. Rev. Biochem. (2025) doi:10.1146/annurev-biochem-030122-043917.

26. Morris, S. K., Baird, C. L. & Lindsley, J. E. Steady-state and rapid kinetic analysis of topoisomerase II trapped as the closed-clamp intermediate by ICRF-193. The Journal of biological chemistry 275, 2613–2618 (2000).

27. Yao, Q., Zhu, L., Shi, Z., Banerjee, S. & Chen, C. Topoisomerase-modulated genome-wide DNA supercoiling domains colocalize with nuclear compartments and regulate human gene expression. Nat. Struct. Mol. Biol. 1–14 (2024) doi:10.1038/s41594-024-01377-5.

28. Mazzocca, M. et al. Chromatin organization drives the search mechanism of nuclear factors. Nat. Commun. 14, 6433 (2023).

29. Roca, J., Ishida, R., Berger, J. M., Andoh, T. & Wang, J. C. Antitumor bisdioxopiperazines inhibit yeast DNA topoisomerase II by trapping the enzyme in the form of a closed protein clamp. Proc. Natl. Acad. Sci. 91, 1781–1785 (1994).

30. Persson, F., Lindén, M., Unoson, C. & Elf, J. Extracting intracellular diffusive states and transition rates from single-molecule tracking data. Nat Methods 10, 265–269 (2013).

31. Mazza, D., Abernathy, A., Golob, N., Morisaki, T. & McNally, J. G. A benchmark for chromatin binding measurements in live cells. Nucleic Acids Res 40, e119 (2012).

32. Seol, Y., Gentry, A. C., Osheroff, N. & Neuman, K. C. Chiral discrimination and writhe-dependent relaxation mechanism of human topoisomerase IIα. The Journal of biological chemistry 288, 13695–13703 (2013).

33. Lee, J. et al. Chromatinization modulates topoisomerase II processivity. Nat. Commun. 14, 6844 (2023).

34. Muñoz-Gil, G., et al. Stochastic particle unbinding modulates growth dynamics and size of transcription factor condensates in living cells. Proc. Natl. Acad. Sci. 119, e2200667119 (2022).

35. Mazzocca, M., Fillot, T., Loffreda, A., Gnani, D. & Mazza, D. The needle and the haystack: single molecule tracking to probe the transcription factor search in eukaryotes. Biochem Soc T 49, 1121–1132 (2021).

36. McKittrick, E., Gafken, P. R., Ahmad, K. & Henikoff, S. Histone H3.3 is enriched in covalent modifications associated with active chromatin. Proc. Natl. Acad. Sci. 101, 1525–1530 (2004).

37. Nguyen, V. Q. et al. Spatiotemporal coordination of transcription preinitiation complex assembly in live cells. Mol Cell 81, 3560–3575.e6 (2021).

38. Muhar, M. et al. SLAM-seq defines direct gene-regulatory functions of the BRD4-MYC axis. Science (New York, N.Y.) 360, 800–805 (2018).

39. Erickson, B., Sheridan, R. M., Cortazar, M. & Bentley, D. L. Dynamic turnover of paused Pol II complexes at human promoters. Genes Dev. 32, 1215–1225 (2018).

40. Mittag, T. & Pappu, R. V. A conceptual framework for understanding phase separation and addressing open questions and challenges. Mol Cell 82, 2201–2214 (2022).

41. McSwiggen, D. T., Mir, M., Darzacq, X. & Tjian, R. Evaluating phase separation in live cells: diagnosis, caveats, and functional consequences. Gene Dev 33, 1619–1634 (2019).

42. Patel, A. et al. A Liquid-to-Solid Phase Transition of the ALS Protein FUS Accelerated by Disease Mutation. Cell 162, 1066–1077 (2015).

43. Singh, V., Johansson, P., Lin, Y.-L., Hammarsten, O. & Westerlund, F. Shining light on single-strand lesions caused by the chemotherapy drug bleomycin. DNA Repair 105, 103153 (2021).

44. Kuzin, V., Wiegard, A., Cameron, D. P. & Baranello, L. TOP1 CAD-seq: A protocol to map catalytically engaged topoisomerase 1 in human cells. Star Protoc 3, 101581 (2022).

45. Pommier, Y. Drugging topoisomerases: lessons and challenges. ACS chemical biology 8, 82–95 (2013).

46. Sciascia, N. et al. Suppressing proteasome mediated processing of topoisomerase II DNA-protein complexes preserves genome integrity. Elife 9, e53447 (2020).

47. Gittens, W. H. et al. A nucleotide resolution map of Top2-linked DNA breaks in the yeast and human genome. Nat Commun 10, 4846 (2019).

48. Heck, M. M., Hittelman, W. N. & Earnshaw, W. C. Differential expression of DNA topoisomerases I and II during the eukaryotic cell cycle. Proceedings of the National Academy of Sciences of the United States of America 85, 1086–1090 (1988).

49. Kimura, K., Saijo, M., Ui, M. & Enomoto, T. Growth state- and cell cycle-dependent fluctuation in the expression of two forms of DNA topoisomerase II and possible specific modification of the higher molecular weight form in the M phase. J. Biol. Chem. 269, 1173–1176 (1994).

50. Earnshaw, W. C., Halligan, B., Cooke, C. A., Heck, M. M. & Liu, L. F. Topoisomerase II is a structural component of mitotic chromosome scaffolds. J. Cell Biol. 100, 1706–1715 (1985).

51. Samejima, K. et al. Mitotic chromosomes are compacted laterally by KIF4 and condensin and axially by topoisomerase IIα. J. Cell Biol. 199, 755–770 (2012).

52. Hildebrand, E. M. et al. Mitotic chromosomes are self-entangled and disentangle through a topoisomerase-II-dependent two-stage exit from mitosis. Mol. Cell 84, 1422–1441.e14 (2024).

53. McPherson, J. P. & Goldenberg, G. J. Induction of apoptosis by deregulated expression of DNA topoisomerase IIalpha. Cancer research 58, 4519–4524 (1998).

54. Yasuda, K., Kato, Y., Ikeda, S. & Kawano, S. Regulation of catalytic activity and nucleolar localization of rat DNA topoisomerase IIα through its C-terminal domain. Genes Genet Syst 95, 291–302 (2020).

55. Frottin, F. et al. The nucleolus functions as a phase-separated protein quality control compartment. Science (New York, N.Y.) 365, 342–347 (2019).

56. Liu, Y. et al. The nucleolus functions as the compartment for histone H2B protein degradation. iScience 24, 102256 (2021).

57. Bülow, S. von, Siggel, M., Linke, M. & Hummer, G. Dynamic cluster formation determines viscosity and diffusion in dense protein solutions. Proc. Natl. Acad. Sci. 116, 9843–9852 (2019).

58. Yu, I. et al. Biomolecular interactions modulate macromolecular structure and dynamics in atomistic model of a bacterial cytoplasm. eLife 5, e19274 (2016).

59. Cho, N. H. et al. OpenCell: Endogenous tagging for the cartography of human cellular organization. Sci. (N. York, NY) 375, eabi6983 (2022).

60. McSwiggen, D. T. et al. A high-throughput platform for single-molecule tracking identifies drug interaction and cellular mechanisms. eLife 12, RP93183 (2025).

61. Das, S. K. et al. Excessive MYC-topoisome activity triggers acute DNA damage, MYC degradation, and replacement by a p53-topoisome. Mol. Cell 84, 4059–4078.e10 (2024).

62. Pomp, W., Meeussen, J. V. W. & Lenstra, T. L. Transcription factor exchange enables prolonged transcriptional bursts. Mol. Cell 84, 1036–1048.e9 (2024).

63. Dessard, M., Manneville, J.-B. & Berret, J.-F. Cytoplasmic viscosity is a potential biomarker for metastatic breast cancer cells. Nanoscale Adv. 6, 1727–1738 (2024).

64. Gal, N. & Weihs, D. Intracellular Mechanics and Activity of Breast Cancer Cells Correlate with Metastatic Potential. Cell Biochem. Biophys. 63, 199–209 (2012).

65. Kalkat, M. et al. MYC Protein Interactome Profiling Reveals Functionally Distinct Regions that Cooperate to Drive Tumorigenesis. Molecular cell 72, 836–848.e7 (2018).

66. Cowling, V. H. & Cole, M. D. The Myc Transactivation Domain Promotes Global Phosphorylation of the RNA Polymerase II Carboxy-Terminal Domain Independently of Direct DNA Binding. Mol. Cell. Biol. 27, 2059–2073 (2007).

67. Dominguez-Sola, D. et al. Non-transcriptional control of DNA replication by c-Myc. Nature 448, 445–451 (2007).

68. Pellanda, P. et al. Integrated requirement of non-specific and sequence-specific DNA binding in Myc-driven transcription. Embo J 40, e105464 (2021).

69. Wiegard, A. et al. Topoisomerase 1 activity during mitotic transcription favors the transition from mitosis to G1. Mol. Cell 81, 5007–5024.e9 (2021).

70. Keppler, A. et al. A general method for the covalent labeling of fusion proteins with small molecules in vivo. Nat. Biotechnol. 21, 86–89 (2003).

71. Lee, J. H., Wendorff, T. J. & Berger, J. M. Resveratrol: A novel type of topoisomerase II inhibitor. Journal of Biological Chemistry 292, 21011–21022 (2017).

72. Broeck, A. V. et al. Structural basis for allosteric regulation of Human Topoisomerase IIα. Nat Commun 12, 2962 (2021).

73. Singh, V. et al. Quantifying DNA damage induced by ionizing radiation and hyperthermia using single DNA molecule imaging. Transl Oncol 13, 100822 (2020).

74. Öz, R. et al. Dynamics of Ku and bacterial non-homologous end-joining characterized using single DNA molecule analysis. Nucleic Acids Res. 49, 2629–2641 (2021).

75. Ershov, D. et al. TrackMate 7: integrating state-of-the-art segmentation algorithms into tracking pipelines. Nat Methods 19, 829–832 (2022).

76. Ewels, P., Magnusson, M., Lundin, S. & Käller, M. MultiQC: summarize analysis results for multiple tools and samples in a single report. *Bioinformatics (Oxford*, England) 32, 3047–3048 (2016).

77. Langmead, B. & Salzberg, S. L. Fast gapped-read alignment with Bowtie 2. Nature methods 9, 357–359 (2012).

78. Li, H. & Durbin, R. Fast and accurate short read alignment with Burrows-Wheeler transform. Bioinformatics (Oxford, England) 25, 1754–1760 (2009).

79. Ramírez, F., Dündar, F., Diehl, S., Grüning, B. A. & Manke, T. deepTools: a flexible platform for exploring deep-sequencing data. Nucleic acids research 42, W187–91 (2014).

80. Flicek, P., et al. Ensembl 2013. Nucleic Acids Res 41, D48–D55 (2013).

